# Histone demethylase Lsd1 is required for the differentiation of neural cells in the cnidarian *Nematostella vectensis*

**DOI:** 10.1101/2020.09.07.285577

**Authors:** James M Gahan, Ivan U. Kouzel, Fabian Rentzsch

## Abstract

The evolution of multicellularity was accompanied by the emergence of processes to regulate cell fate, identity and differentiation in a robust and faithful manner. Chromatin regulation has emerged as a key process in development and yet its contribution to the evolution of such processes is largely unexplored. Chromatin is regulated by a diverse set of proteins, which themselves are tightly regulated in order to play cell/ tissue-specific functions. Using the cnidarian *Nematostella vectensis*, a model for basal metazoans, we explore the function of one such chromatin regulator, Lysine specific demethylase 1 (Lsd1). We generated an endogenously tagged allele and show that the expression of NvLsd1 is developmentally regulated and higher in differentiated neural cells than their progenitors. We further show, using a CRISPR/Cas9 generated mutant that loss of *NvLsd1* leads to several distinct developmental abnormalities. Strikingly, *NvLsd1* loss leads to the almost complete loss of differentiated cnidocytes, cnidarian-specific neural cells, which we show to be the result of a cell-autonomous requirement for *NvLsd1*. Together this suggests that complex regulation of developmental processes by chromatin modifying proteins predates the split of the cnidarian and bilaterian lineages, approximately 600 million years ago, and may constitute an ancient feature of animal development.

## Introduction

The evolution of animals and other multicellular organisms was accompanied by the emergence of the processes of cell type specification and differentiation. These processes are tightly controlled in order to produce adults containing, in some cases, billions of cells and hundreds of different cells types originating from a single celled embryo. The regulation of cell differentiation is largely governed on the transcriptional level. This has classically been viewed as the purview of DNA binding transcription factors that regulate sets of genes in a spatial and temporally restricted manner to drive cell identity and differentiation (1). In more recent years the role of chromatin modifications in this process has become more appreciated (2-4). In particular, it is now clear that chromatin modifiers are not static, permissive actors in the process of cell differentiation but rather play an active role (4). Indeed, chromatin modifiers such as those involved in histone lysine methylation and demethylation often play cell or tissue specific roles (5, 6). Such chromatin modifiers can themselves be regulated, at the level of expression, splicing, localization and activity in order to allow for differential functions in different cells or tissues. Transcriptional regulation by chromatin modifiers is present in unicellular eukaryotes, but it is currently not understood at what point in the evolution of animals this function became integrated into the regulatory programs that control the differentiation of specific cell types. Here, we address this question by analyzing the role of the conserved chromatin regulator Lsd1/KDM1A in the development of an early-branching metazoan, the sea anemone *Nematostella vectensis*.

Lsd1 acts, in the majority of cases, to remove mono- or di-methylation on lysine 4 of histone H3 (H3K4me1/me2) (7). It does so in a complex with Histone Deacetylase 1/2 (HDAC1/2) and Corepressor of REST (CoREST) in order to repress target genes (8-14). In other cases, however it has been shown that Lsd1 is involved in demethylation of H3K9 or H4K20 and can act as a transcriptional co-activator (15-18). Loss of Lsd1 in all cells leads to early lethality in mice (15), whereas tissue specific manipulations of Lsd1 function revealed roles in the development and homeostasis of several organs and cell types (15, 19-27). Lsd1 has been shown to be particularly important in the nervous system where it regulates neurogenesis at several levels. In rodents, Lsd1 levels are higher in neural stem/progenitor cells and decrease as differentiation progresses (28-30). Lsd1 plays roles in the maintenance of progenitor identity and proliferation and the decrease in Lsd1 levels is required for neural differentiation (28-30). Lsd1 is also important in the later stages of neural differentiation, for example for cortical migration of differentiating neurons in mice (31) and for the terminal differentiation of rod photoreceptors (32). In humans, LSD1 mutations have been linked to a developmental disorder that includes severe cognitive impairment (33, 34). In contrast to rodents, however, LSD1 levels remain largely unchanged or increase slightly throughout differentiation in the human nervous system (18, 35). In *Drosophila melanogaster,* loss of Lsd1 leads to a range of developmental abnormalities without being lethal (36). In *Caenorhabditis elegans,* the Lsd1 homolog SPR5 is required to erase H3K4me2 in primordial germ cells and its loss leads to a transgenerational loss of fertility (37).

Despite the importance of chromatin regulation during development, it is only well understood in a small number of “model” systems. All of these models belong to the Bilateria, the clade of animals containing most animal phyla (Figure 1A) In order to understand how these processes evolved it is important to look at other, more early diverging groups of animals. *Nematostella vectensis*, the starlet sea anemone, belongs to the sister group to the Bilateria, the Cnidaria (Figure 1A), which separated from the lineage leading to bilaterians approximately 600 million years ago (38, 39). This key phylogenetic position along with the expanding molecular and genetic toolkit available, make *Nematostella* a useful system in which to address developmental questions in detail while also revealing evolutionary aspects of developmental processes (40). Descriptive work at the whole organism-level in *Nematostella* has shown that active chromatin modifications display similar patterns genome wide to those seen in other animals (41) and studies in other cnidarians suggest roles for histone acetylation in development (42, 43).

**Figure 1:**
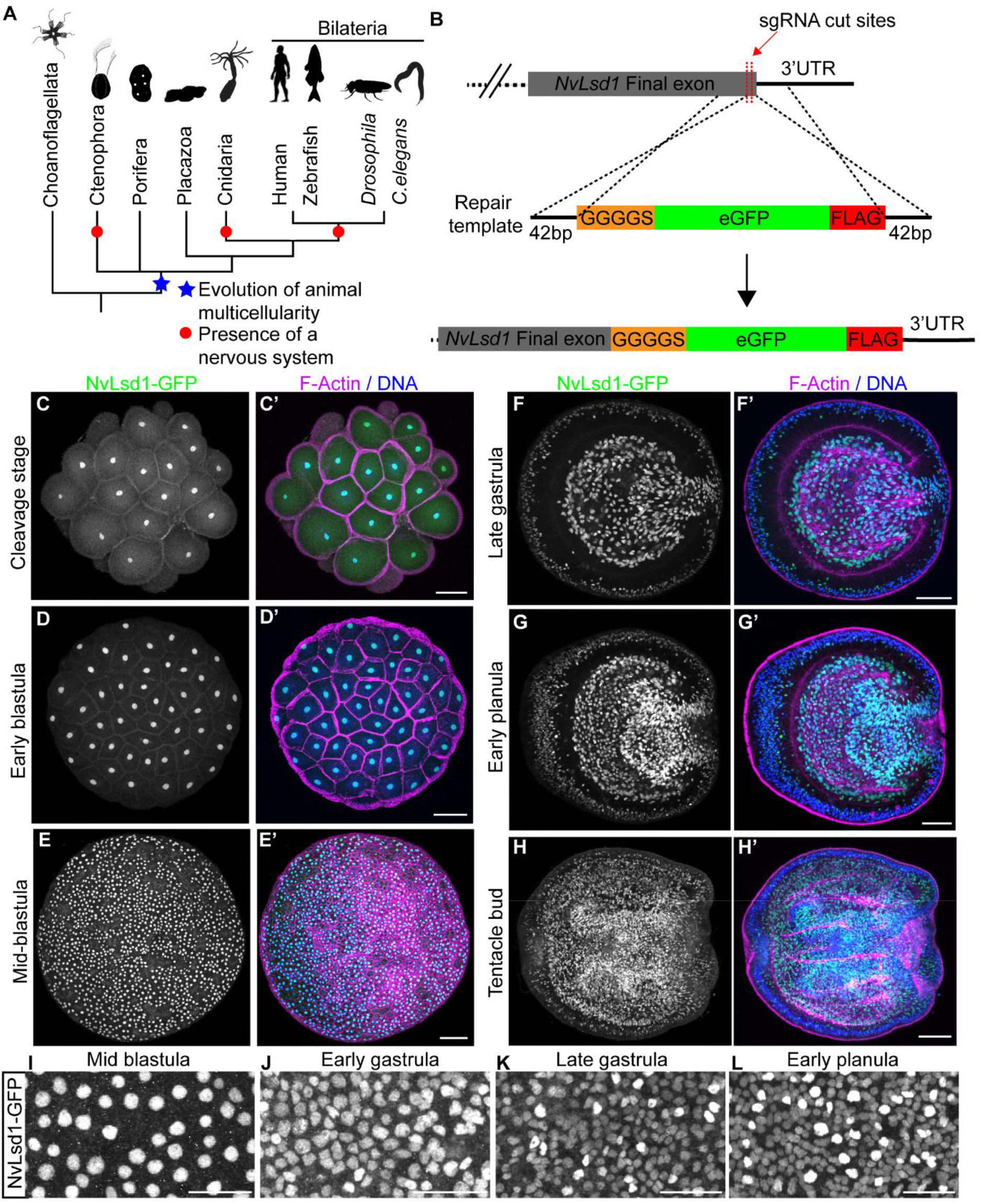
NvLsd1 is ubiquitously expressed but developmentally regulated. (**A**) Reduced phylogeny showing the position of the 4 non-bilaterian animal lineages with respect to the Bilateria. The evolution of animal multicellularity is annotated by a blue star and the lineages possessing a nervous system are marked with a red circle. Phylogeny based on (39). (**B**) Schematic showing the strategy for generating the *NvLsd1^GFP^* knock-in allele by CRISPR-Cas9 mediated homologous recombination. (**C-H**) Confocal images of immunofluorescence staining on *NvLsd1^GFP^* animals. Stage is shown on the left. Oral is shown to the right in F-H. NvLsd1-GFP is shown in green, DNA in blue and F-Actin in magenta. **F-H** show mid-lateral views. (**I-L**) Close ups of NvLsd1-GFP in the ectoderm at different stages. Scale bars: 50 μm (C-H), 20 μm (I-L).

*Nematostella* development proceeds via a hollow blastula stage and gastrulation by invagination to form a free-swimming planula larva. About one week after fertilization, the planula settles and develops into a sessile primary polyp that uses a ring of tentacles surrounding the only body opening to catch prey (44-46) (Figure S1A). As for all cnidarians, *Nematostella* tissue has only two germ layers, which we here call ectoderm and endomesoderm (orange in Figure S1A). The *Nematostella* nervous system consists of a nerve net, thought to be an ancestral characteristic of animal nervous systems (47). As in other cnidarians, there are three major, morphological cell types in the *Nematostella* nervous system; sensory cells, ganglionic neurons and cnidocytes; cnidarian specific neural cells also known as stinging cells (48, 49). Recent years have seen advances in the understanding of the development of this nervous system, with a focus on transcription factors and signaling pathways (50-62). How chromatin is regulated during neurogenesis and how chromatin modifiers are integrated with cell differentiation in *Nematostella* is unknown.

Here we use Lsd1 as a paradigm for a developmentally regulated chromatin modifier to examine chromatin based regulation of development and cell differentiation in *Nematostella*. Employing CRISPR/Cas9-mediated genome editing, we generated an endogenously tagged allele and show that NvLsd1 is expressed at elevated levels in differentiated neural cells. Morphological and molecular analyses of a loss-of-function mutant identify defects in the differentiation of cnidocytes that can be rescued by cnidocyte-specific re-expression of *NvLsd1*. These results suggest that *NvLsd1* has an ancient function in neural cell differentiation.

## Results

### NvLsd1 is ubiquitously expressed but developmentally regulated

*Nematostella* has a single, highly conserved homolog of Lsd1 (63), hereafter called *NvLsd1* (Figure S2). RNA *in-situ* hybridization showed that at early gastrula, *NvLsd1* is expressed throughout the embryo but more highly in scattered single cells in the ectoderm (Figure S1B). In late gastrula and planula stages, this pattern is maintained and strong expression is seen in the pharynx and endomesoderm (Figure S1C, D). Since Lsd1 levels are subject to posttranscriptional regulation in other animals, we generated an endogenous allele tagged with eGFP and a FLAG tag using CRISPR-Cas9 mediated homologous recombination in order to better visualize NvLsd1 expression (Figure 1B). From here on, we refer to the animals carrying this allele as *NvLsd1^GFP^* animals. We confirmed the insertion of a single copy of GFP using PCR across the insertion site from gDNA (Figure S3A, D) and sequencing of this band. We also generated cDNA from *NvLsd1^GFP^* animals and were able to clone a full-length cDNA containing the full *NvLsd1* coding sequence, the inserted sequences as well as an intact 3’ UTR (Figure S3B, C). In addition, we performed a western blot using an antibody against GFP and found a single band corresponding to the approximate size of the NvLsd1-GFP fusion protein (Figure S3E). Analysis of *NvLsd1^GFP^* animals revealed that NvLsd1 is a ubiquitous protein found in all nuclei throughout development (Figure 1C-L) except on mitotic chromatin, which is devoid of NvLsd1-GFP signal (Figure S4A), similar to what was previously shown in mammalian cells (64). As seen by RNA *in-situ* hybridization, high levels of NvLsd1-GFP expression are seen in the pharynx and endomesoderm (Figure 1F-H). In addition, there are indeed two populations of nuclei in the ectoderm; one with low and one with high levels of NvLsd1-GFP, from here on referred to as NvLsd1^low^ and NvLsd1^high^ cells (Figure 1I-L). These two populations can also be distinguished by flow cytometry (Figure 2F and S5A). This heterogeneity in NvLsd1-GFP levels becomes more distinct as development progresses (Figure 1I-L). Together this shows that NvLsd1 is ubiquitously expressed but that its levels are regulated in a cell-type-specific manner during development.

**Figure 2:**
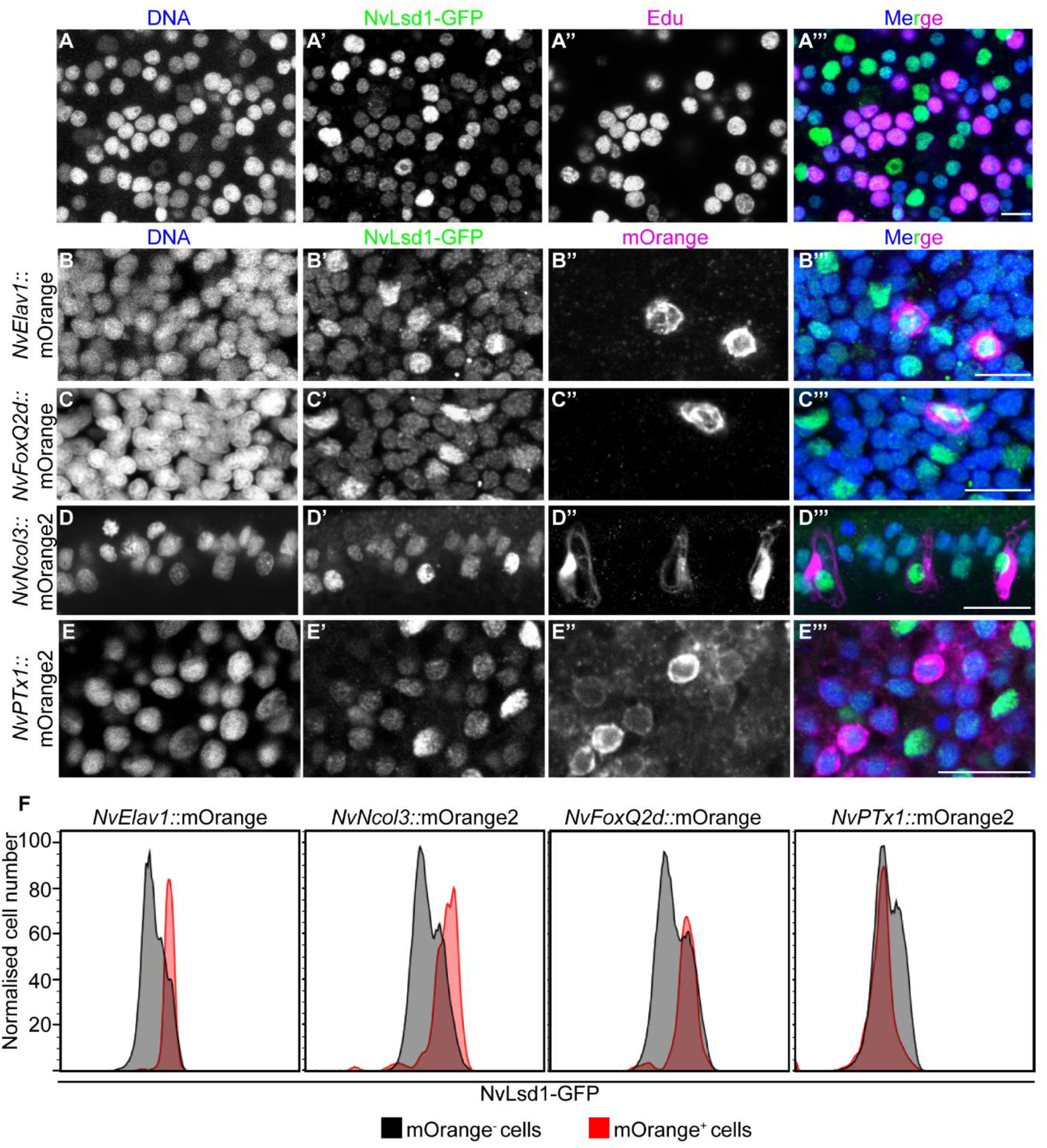
NvLsd1 is highly expressed in differentiated neural cells. (**A**) Confocal images showing close up of the ectoderm of *NvLsd1^GFP^* late gastrula treated with Edu for 4 hours stained with anti-GFP antibody (green) and Click-IT Edu (Magenta). DNA is shown in blue. (**B-E**) Confocal images of immunofluorescence staining on late planula of *NvLsd1^GFP^* animals crossed with *NvElav1*::mOrange (**B**), *NvFoxQ2d*::mOrange (**C**), *NvNcol3*::mOrange2 (**D**) and mid-planula crossed with *NvPTx1*::mOrange2 (**E**) transgenics. DNA is shown in blue, GFP in green and mOrange in magenta. (**F**) Histograms of flow cytometry data on Lsd^*GFP*^ animals crossed to the indicated transgenic lines. The X-axis shows the GFP fluoresce levels (units are arbitrary) and the Y-axis shows the cell number normalized to mode to correct for the different numbers of cells in the mOrange^+^ and mOrange^−^ gates. Scale bars: 20 μm.

### NvLsd1 levels are high in differentiated neural cells

In order to determine the identity of the NvLsd1^low^ and NvLsd1^high^ cells we first performed EdU labelling experiments to mark proliferating cells. At late gastrula stage we labelled with EdU for 30 minutes or four hours. 30 minutes incubation labels mostly S phase cells while four hours incubation leads to labeling of all cell in S-phase, G2 and likely early G1. This can be seen as after four hours of EdU incubation all mitotic nuclei are EdU positive (Figure S4A). We performed this staining in *NvLsd1^GFP^* animals and could see that in all cases NvLsd1^high^ cells were EdU negative (Figure 2A and S3B). This was also true later at mid-planula stage (Figure S4C). Given the known role of Lsd1 in the mammalian nervous system we next investigated NvLsd1-GFP levels in different neural cell types. To do this we crossed *NvLsd1^GFP^* animals to previously published neural reporter lines: *NvFoxQ2d*::mOrange which labels a population of sensory cells (53), *NvElav1:*:mOrange which labels a non-overlapping population of sensory and ganglion cells (53, 62) and *NvNcol3:*:mOrange2 which labels cnidocytes (65). In all cases, the mOrange^+^ cells have high levels of NvLsd1-GFP at late planula stage (Figure 2B-D). In contrast, ectodermal gland cells labelled in the *NvPTx1*::mOrange2 line (66) do not overlap with the NvLsd1^high^ population (Figure 2E) at mid-planula stage, indicating that high levels of NvLsd1 are not a general feature of differentiated cells. To quantify this we performed flow cytometry at the same stages. In the case of each transgenic line, we could compare the levels of NvLsd1-GFP in mOrange^+^ versus mOrange^−^ cells (Figure S5A). This revealed that in the *NvElav1:*:mOrange, *NvFoxQ2d*::mOrange and *NvNcol3:*:mOrange2 line the mOrange^+^ cells fall largely within the NvLsd1^high^ population while in the *NvPTx1*::mOrange2 line far fewer mOrange^+^ cells are NvLsd1^high^ (Figure 2F and S5B). Together this shows that NvLsd1 levels are low in proliferating cells and are higher in differentiated neural cells.

### Loss of *NvLsd1* is not lethal but leads to developmental abnormalities

To analyze the function of *NvLsd1* we generated a mutant allele using CRISPR-Cas9. We targeted an exon containing a lysine at position 644 or K644 (K661 in humans) which is important for catalytic function and is located in a highly conserved portion of the protein (Figure S2). This residue interacts with the cofactor FAD and has been shown to be required for catalytic activity *in vitro* (8, 10, 11, 67), although mutation of it does not completely abolish enzymatic activity (68). We generated a mutant line containing a 4 base pair deletion upstream of this lysine leading to a truncated protein missing a large portion of the amine oxidase domain (Figure 3A). This mutation is similar to a zebrafish mutant that has been shown to be a null allele (69). In order to trace the mutation we crossed mutants to *NvLsd1^GFP^* animals generating animals with one GFP tagged allele and one mutant allele, *NvLsd1^GFP/-^*. In *NvLsd1^GFP/-^ x NvLsd1^GFP/-^* crosses, 75% of the animals have at least one copy of the *NvLsd1^GFP^* allele, hereafter called control, and the homozygous mutant animals are GFP negative, hereafter *NvLsd^−/−^* (Figure 3B). This allowed us to distinguish *NvLsd^−/−^* animals from control by absence or presence of GFP signal, respectively. We confirmed the efficacy of this approach via sequencing (Figure S6A). *NvLsd^−/−^* animals develop normally until late planula (Figure 3F, G). We counted the number of animals at late planula stage and show approximately Mendelian ratios of *NvLsd^−/−^* animals and controls, indicating no major defects in survival (Figure 3C). We also quantified the percentage of animals that developed into primary polyps and saw no difference between mutants and controls (Figure 3D). *NvLsd^−/−^* primary polyps are shorter than controls and have shorter tentacles but otherwise appear morphologically normal (Figure 3E, H and I). A more detailed morphological examination using immunofluorescence also revealed no obvious morphological differences at late planula stage (Figure 3J, K). At primary polyp stage, although *NvLsd^−/−^* animals are shorter, all major morphological structures have formed, i.e. animals have four tentacles, a pharynx, and the internal tissue folds known as mesenteries (Figure 3L, M). These animals fail to begin feeding normally and never develop fully (Figure S6B). The mutation does not, however, appear to be lethal and these animals can survive for greater than 3 months although they do not grow to maturity (Figure S6B). Together this shows that, although *NvLsd1* loss is not lethal during embryonic stages, *NvLsd^−/−^* animals have altered development and do not survive until adulthood.

**Figure 3:**
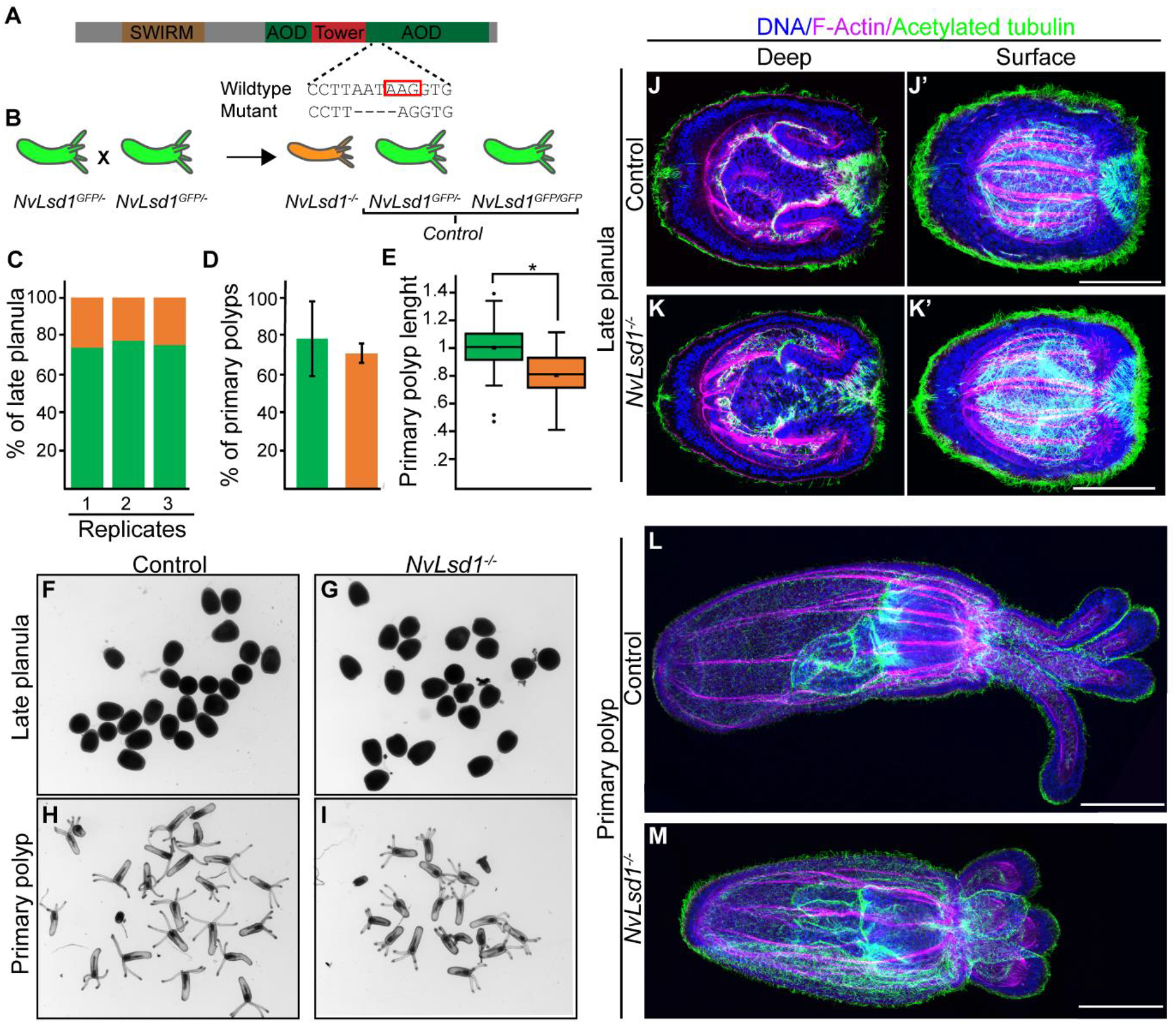
Generation and characterization of *NvLsd1* mutants. (**A**) Schematic showing the structure of the *NvLsd1* protein and the position and sequence of the mutation. The red box indicates the position of K644. (**B**) Cartoon depicting the crossing scheme used in all experiments with the *NvLsd1* mutants. (**C**) Quantification of the percentage of late planula which were control (Green) or *NvLsd1^−/−^* (Orange) across 3 independent biological replicates. (**D**) Percentage of control and *NvLsd1^−/^* late planula which develop into primary polyps. Bar chart shows the mean +/− standard deviation from 3 biological replicates. (**E**) Measurement of body column length in control and *NvLsd1^−/−^* primary polyps. The total body length is normalized to control. Data is combined from 3 independent biological replicates. Statistical significance was determined using the Student’s T-test * p<1e-7. (**F-I**) Brightfield pictures of control or *NvLsd1^−/−^* animals at late planula and primary polyp stage. Genotype is shown on top and stage is shown to the left. (**J-M**) Confocal images of immunofluorescence staining on control and *NvLsd1^−/−^* late planula (**J,K**) and primary polyp (**L-M**). DNA is shown in blue, acetylated tubulin in green and F-actin in magenta. J and K show several projected confocal slices from the inside of the planula and J’ and K’ show several projected slices from closer to the surface. Scale bar: 50 μm.

### Transcriptomic analysis of *NvLsd1* mutants reveals a possible role for *NvLsd1* in cnidocyte differentiation

Next, we wanted to analyze the transcriptional changes occurring due to mutation of *NvLsd1*. To do this, we performed RNAseq on *NvLsd1* mutants and controls (consisting of *NvLsd1^GFP/-^* and *NvLsd1^GFP/GFP^ animals*) in quadruplicate at two different developmental stages: 4 day-old late planula and 13 day-old primary polyp (Figure 4A). Principal component analysis (PCA) showed that, as expected, the majority of variance separates samples based on developmental stage (the first principal component, PC1). However, the second principal component (PC2) separates animals based on whether they are *NvLsd^−/−^* or control (Figure 4B). Differential gene expression analysis revealed 1641 differentially expressed genes in late planula (580 up-regulated and 1061 down-regulated) and 7080 in primary polyp (3439 up-regulated and 3641 down-regulated) (Figure 4C, D). The lower number of genes differentially expressed at late planula stage correlates with the less severe phenotype seen at this stage (see also later sections) but it is also important to note that maternal *NvLsd1* persists late in development and could be masking the effects of zygotic loss of *NvLsd1* (Figure S6C, D). Comparing genes differentially expressed at the different stages revealed that 1158 out of 1641 genes (**≈** 71%) differentially expressed in late planula are also differentially expressed in primary polyps (Figure 4E). *NvLsd1* itself was among the downregulated genes in both primary polyp and late planula samples (Figure S7A). Given the high levels of NvLsd1 in differentiated neural cells, we compared the differentially expressed genes to a previously published transcriptome from *NvElav1*::mOrange^+^ neurons (52) which we have re-analyzed here and to the transcriptome of *NvNcol3*::mOrange2^+^ cnidocytes also produced here. Importantly, both these transcriptomes were also generated at 13 day primary polyp. We observed only limited overlap of genes up-regulated in mutants with genes up-regulated in either cell population (~16-23%) (Figure S7B, C). However, when we looked at genes down-regulated in mutants, we see that for both late planula and primary polyp there is a larger overlap with genes up-regulated in cnidocytes (~67% and ~31%, respectively) (Figure 4F) but not with genes up-regulated in neurons (~12% and ~13% respectively) (Figure S7D). These include genes known to be expressed in cnidocytes such as NEP6 and NEP16 (70) (Figure S7A). To test if these overlaps were statistically significant, we used the GeneOverlap R package (Shen and Sinai, 2020) which implements Fisher’s exact test to calculate if the overlap is significant and provides several metrics, such as p-value (significance) and Odds ratio (strength of association). An Odds ratio of 1 or less indicates no association while an Odds ratio above 1 indicates an association between the gene sets and the higher the number, the greater the association. The overlap between genes down-regulated in *NvLsd1* mutants and *NvNcol3*::mOrange2^+^ cnidocytes was significant while the overlap with genes up-regulated in *NvElav1*::mOrange^+^ neurons was not (Figure 4G). The overlaps between genes up-regulated in *NvLsd1* mutants and those up-regulated in *NvElav1*::mOrange^+^ were also not significant. The genes up-regulated in primary polyp stage did not overlap significantly with those upregulated in *NvNcol3*::mOrange2^+^ cnidocytes, however, those up-regulated in late planula did have a significant overlap albeit with a relatively low Odds ratio and higher p-value (Figure 4G). We also performed more stringent analysis for the selection of differentially expressed genes by raising the log2 fold change threshold from 0 to 1, and hence, getting the genes with more substantial changes due to mutation of *NvLsd1* (standard DESeq2 approach). We found the same results although we now see an overlap between genes up-regulated in primary polyp and *NvNcol3*::mOrange2^+^ cells although with a very high p-value and low Odds ratio (Figure S7E). Together this indicates that loss of *NvLsd1* may affect cnidocyte formation.

**Figure 4:**
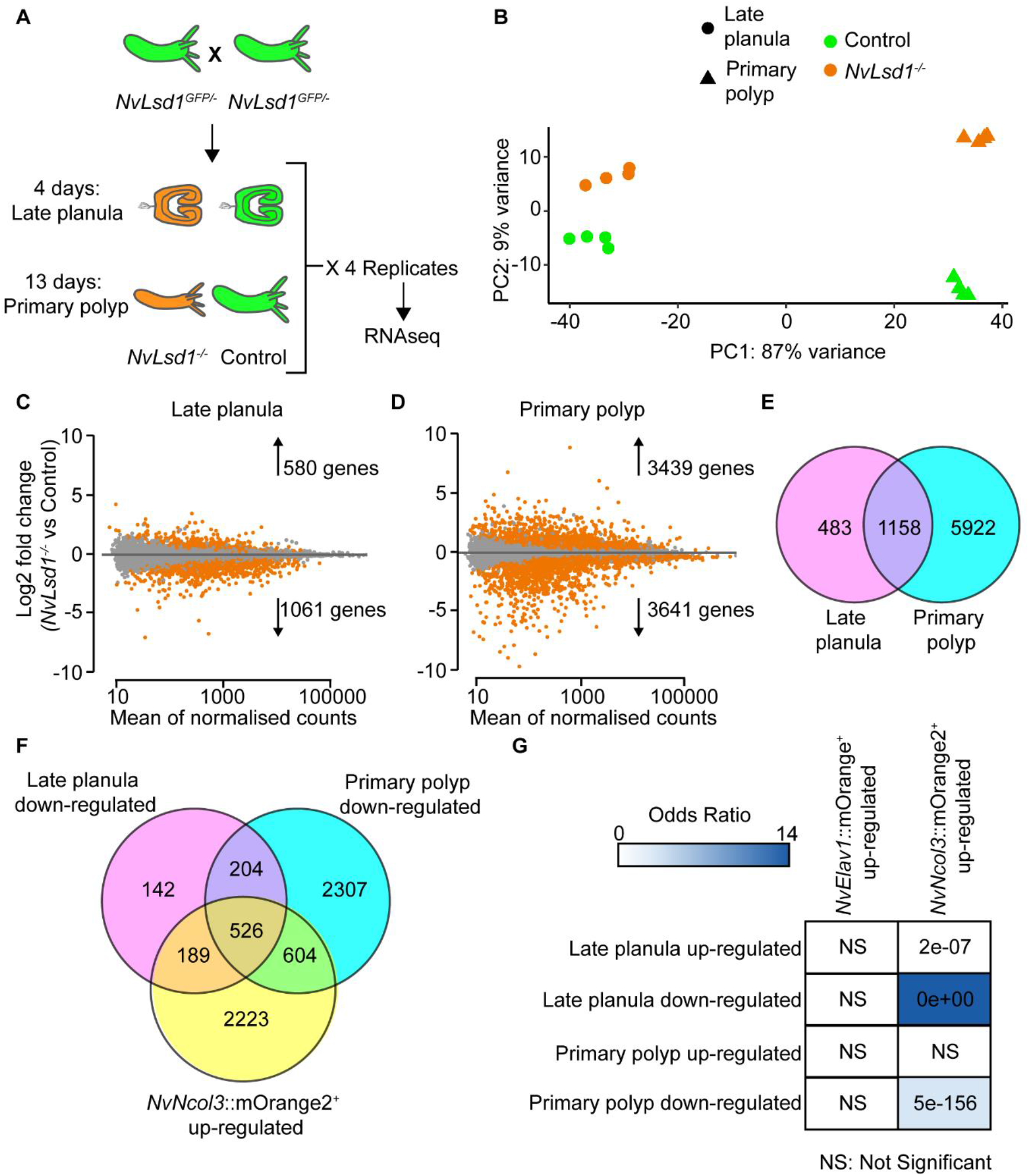
Transcriptomic analysis of Lsd1 mutants. (**A**) Cartoon depicting the experimental design for the transcriptome analysis of *NvLsd1* mutants. Controls are shown in green and mutants are shown in orange. (**B**) PCA plot using the gene counts after variance stabilizing transformation (VST). Late planula samples are shown as circles and primary polyp as triangles. Experimental conditions are encoded with color: green for controls and orange for mutants. (**C-D**) MA-plots of up- and down-regulated genes between *NvLsd1^−/−^* and control in late planula (**C**) and primary polyps (**D**). Differentially expressed genes are shown in orange at significance level of 0.05. (**E**) Venn diagram comparing all differentially expressed genes at late planula and primary polyp stages. (**F**) Venn diagram comparing genes down-regulated in *NvLsd1^−/−^* late planula and primary polyp with genes up-regulated in *NvNcol3*::mOrange2^+^ cells. (**G**) Comparison of the overlap between up- and down-regulated genes in *NvLsd1^−/−^* late planula and primary polyps with those up-regulated in *NvNcol3*::mOrange2^+^ and *NvElav1*::mOrange^+^ cells using the GeneOverlap R package. The strength of the blue color indicates the odds ratio and numbers indicate the p-value, calculated using Fisher’s exact test.

### *NvLsd1* mutants have defects in cnidocyte differentiation but not specification

Having found a significant overlap between downregulated genes in the *NvLsd^−/−^* animals and those upregulated in cnidocytes, we next looked at cnidocyte differentiation in *NvLsd^−/−^* animals. Cnidocytes are cnidarian specific neural cells that are responsible for prey capture and defense. Cnidocytes contain a specific organelle, called the cnidocyst, which forms as a large post-Golgi compartment. This pressurized organelle contains a tubule that is extruded like a harpoon upon explosive rupture of the cnidocyst (71). Different stages in the development of this neural cell type can be visualized by monitoring the formation and maturation of the cnidocyst. An antibody against NvNcol3 labels the forming cnidocyst wall but does not label the fully mature cnidocyst (72). A modified DAPI staining protocol, on the other hand, labels the matrix of the mature cnidocyst (73). Staining at the late planula stage revealed an overall reduction in the number of mature cnidocysts in *NvLsd^−/−^* animals (Figure 5A-C). NvNcol3 staining in controls shows developing cnidocysts with a regular, elongated shape (Figure S5 A’, D). In *NvLsd^−/−^*animals, however, patchy and irregular staining was observed suggesting that although cnidocytes are present, they are not able to complete their differentiation (Figure 5B’, E). At polyp stage there is an even larger reduction in mature cnidocysts (Figure 5F, G) and the same change in the pattern of NcNcol3 staining is observed (Figure 5F-H). Since *NvLsd1* is ubiquitously expressed, we tested whether the role in cnidocyte development was cell autonomous. There are currently no conditional loss-of-function approaches established for *Nematostella* and we therefore decided to use a cell-type-specific rescue approach. We expressed *NvLsd1* under the *NvPOU4* promoter that, in the tentacles, is primarily expressed in post-mitotic cnidocytes (52). We generated F0 mosaic transgenics and stained for cnidocytes using *NvPOU4*::*NvHistoneH2B* as a control. In some cases, we see small patches of expression with few positive nuclei. We exclude these from our analysis as, in uninjected *NvLsd1* mutants, there is always a small number of cnidocytes in the tentacles (Figure 5G) making it difficult to analyze the effects of expression in just a few cells. In larger patches, expression of *NvPOU4*::*NvLsd1* always led to a rescue of the cnidocyte phenotype as seen by corresponding, overlapping patches of DAPI^+^ cnidocysts (n=9/9) (Figure 6A). *NvPOU4*::*NvHistoneH2B* expression, however, never led to such an effect (n=16/16) (Figure 6B). We also generated a mutant version of *NvLsd1* bearing two single amino acid changes that completely block catalytic activity in human LSD1 (68), NvLsd1^K644A/A520E^. Expression of this mutant was unable to rescue the effect of loss of *NvLsd1* in cnidocytes (n=6/6) (Figure 6C). Together this data shows an important role for *NvLsd1* in cnidocyte maturation.

**Figure 5.**
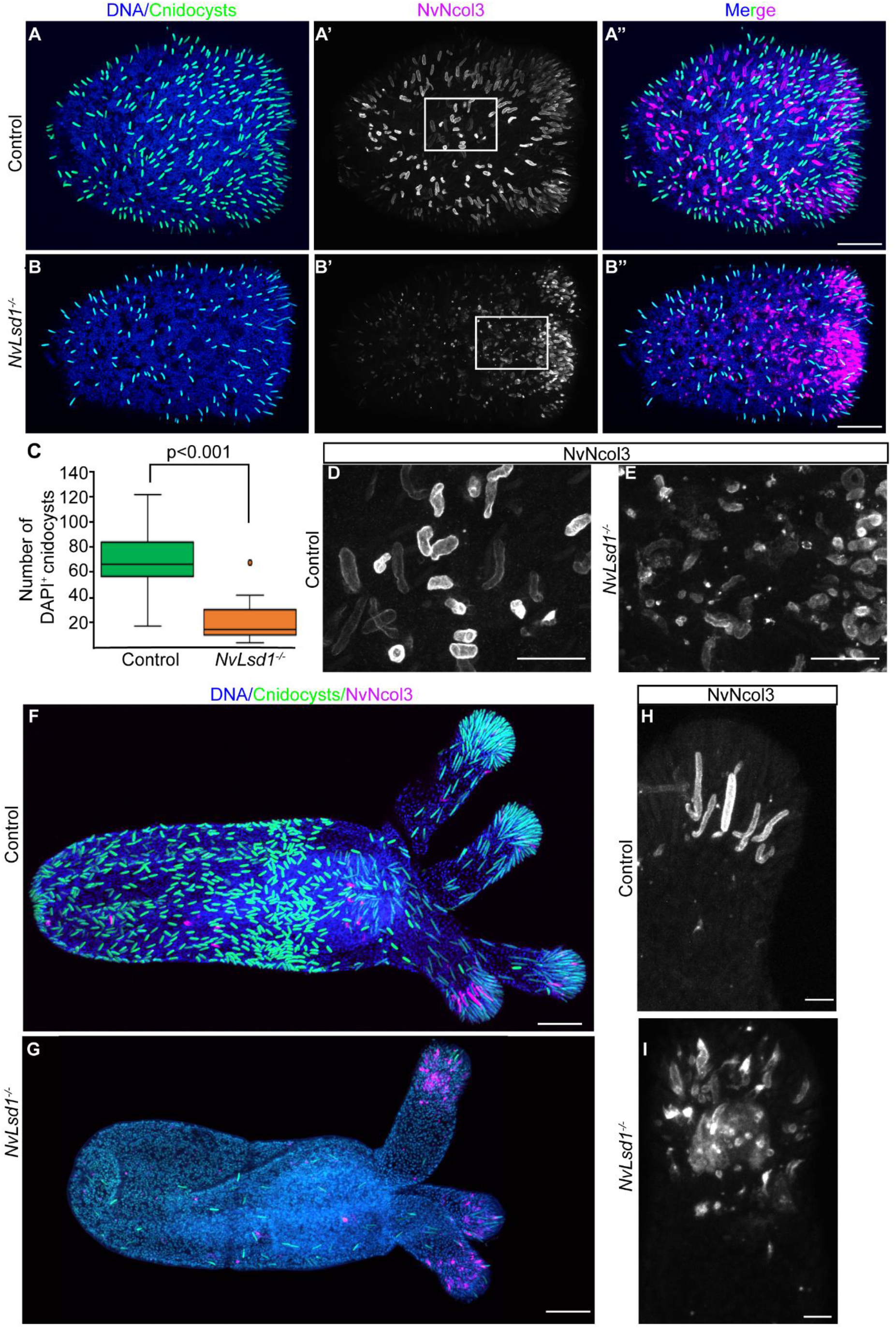
Lsd1 mutants have major defects in cnidocyte differentiation. (**A, B**) Confocal images of immunofluorescence staining on control and *NvLsd1^−/−^* late planula showing DNA in blue, mature cnidocysts in green and NvNcol3 in magenta. (**C**) Quantification of mature DAPI^+^ cnidocysts from control and *NvLsd1^−/−^* late planula. The number of cnidocysts in a 100 μM section in the center of each larva was counted. Statistical significance was calculated using a Student’s t-test. (**D, E**) Close ups of the regions highlighted in A’ (**D**) and B’ (**E**) showing NvNcol3 staining in grey. (**F, G**) Confocal images of Immunofluorescence staining on control and *NvLsd1^−/−^* primary polyps. DNA is shown in blue, mature cnidocysts in green and NvNcol3 in magenta. (**H, I**) High magnification of NvNcol3 staining in the tentacles of primary polyps in F (**H**) and G (**I**) showing NvNcol3 staining in grey. Scale bars: 50 μm (A, B, F and G), 20 μm (D, E) and 10 μm (H, I).

**Figure 6:**
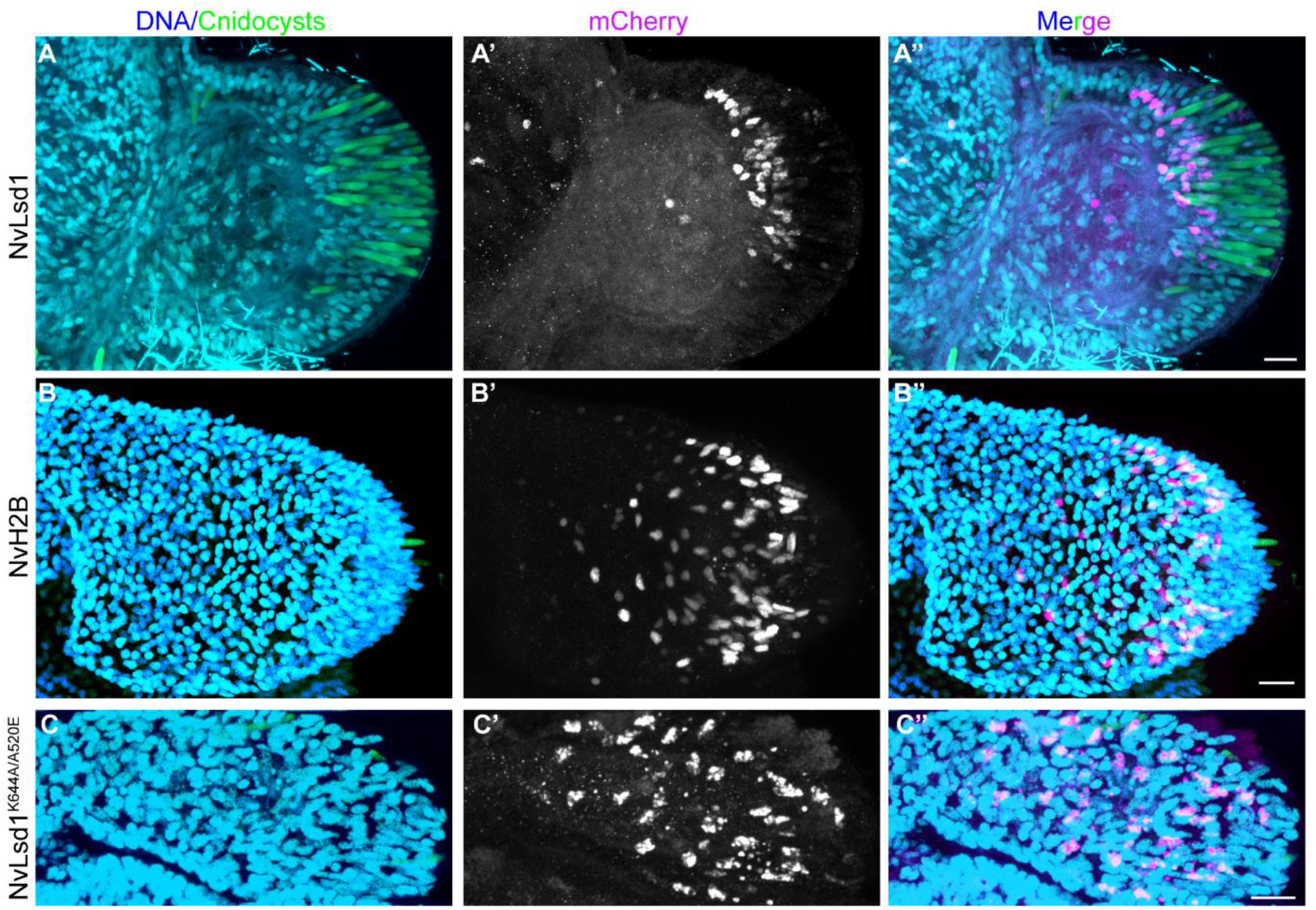
NvLsd1 is required cell autonomously for cnidocyte differentiation. (**A-C**) Immunostaining on NvLsd1^−/−^ animals injected with *NvPOU4*:NvLsd1-mCherry (**A**), *NvPOU4*:NvH2B-mCherry (**B**) or *NvPOU4*:NvLsd1^K644A/A520E^-mCherry (**C**). DNA is shown in blue, DAPI^+^ cnidocysts in green and mCherry in magenta. Scale bars: 10 μm.

### Loss of *NvLsd1* has only minor effects on neurons

Although there was no significant overlap between genes differentially expressed in *NvLsd^−/−^* animals and genes up-regulated in *NvElav1*::mOrange^+^ neurons, we analyzed the effects of loss of *NvLsd1* on these neurons as NvLsd1 levels were shown to also be high in these cells. In primary polyps, the expression of the transgenic reporter is less uniform with many mOrange^+^ punctae, which are not present in controls and with an apparent expansion of fluorescence into endodermal epithelial cells (Figure S8A, B). We also looked at *NvFoxQ2d*::mOrange^+^ neurons but did not see gross changes in these cells in *NvLsd^−/−^* animals compared with controls (Figure S8C, D).

## Discussion

Lsd1 is an important chromatin regulator involved in several steps of the neurogenesis process in mammals. Here we show that Lsd1, in the sea anemone *Nematostella*, is developmentally regulated and plays a crucial role in the differentiation of cnidocytes, a cnidarian specific neural cell type.

The levels of chromatin regulators are often tightly controlled in a cell and tissue specific manner in bilaterians. Lsd1 is a prime example of this where there are many examples of its complex regulation. In *Nematostella,* we have shown that NvLsd1 levels are also regulated during neurogenesis with higher levels seen in differentiated cells. This is similar to observations in human cells (18, 35). This regulation appears to be both transcriptional and post-transcriptional. We see by RNA in-situ hybridization that *NvLsd1* mRNA levels are heterogeneous (Figure S1B) and it is also up-regulated in the *NvElav1*::mOrange^+^ cells indicating regulation on the transcriptional level. We have also shown that some maternal NvLsd1 lasts late into development and we have shown that, although levels are much lower overall, there is still a heterogeneity between cells in the amount of maternal protein present at late planula stage (Figure S6C, D). As this heterogeneity is derived from maternal mRNA and/or maternal Lsd1 protein, it must represent a level of post transcriptional control of NvLsd1 levels. It will be interesting in the future to dissect the transcriptional and post-transcriptional regulation of Lsd1 and compare it to that of mammals to understand if there are conserved processes involved.

In mice, Lsd1 mutants are lethal at early embryonic stages (15) while in zebrafish, *Drosophila* and *C.elegans,* mutants are viable but display several phenotypic abnormalities (36, 37, 69). Here we show that loss of zygotic *NvLsd1* is not lethal in *Nematostella* and in fact, animals survive for many months. It is, however, possible that *NvLsd1* is crucial for early development as NvLsd1 levels are very high maternally and some maternal NvLsd1 lasts late into development. Indeed, in mice maternal Lsd1 is required for the maternal to zygotic transition (74, 75) and in fish it has also been proposed that the large maternal pool of Lsd1 may be the reason for a lack of more severe phenotypes early in development (69). We have not been able to grow *NvLsd1^−/−^* animals to sexual maturity likely due to their inability to feed/grow properly. Therefore, studying this maternal pool of *NvLsd1* will require the establishment of new tools in *Nematostella*.

The function of Lsd1 in the nervous system has so far only been well demonstrated in mammals where Lsd1 is playing many different roles. Here we show that *NvLsd1* also has a role in the nervous system of *Nematostella*. The strongest evidence for this in the requirement for *NvLsd1* for proper cnidocyte differentiation. Cnidocytes, although a cnidarian specific cell type, are bona fide neural cells (48, 49). Loss of *NvLsd1* leads to a loss of mature cells but does not appear to lead to major defects in cell specification. This can be seen by the continued presence of NvNcol3^+^ immature cnidocytes despite the severe reduction in mature cells. *NvNcol3* is also not differentially expressed in mutants. Additionally, transcription factors *NvPaxA* and *NvPou4*, which are known regulators of cnidocyte differentiation (52, 58), are not differentially expressed in either late planula or primary polyps. In contrast, *Cnido-Jun* and *Cnido-Fos1* are downregulated in mutants at both stages (Figure S7A). These two transcription factors have recently also been shown to be required for cnidocyte differentiation (65). This indicates that the effect of loss of *NvLsd1* occurs downstream of *PaxA* and *NvPou4* but upstream of other regulators like *Cnido-Jun* and *Cnido-Fos1*. We do not, however, have a full picture of the gene regulatory network involved in cnidocyte differentiation and it is therefore currently difficult to place *NvLsd1* function within this network. The rescue experiments further showed that expressing *NvLsd1* in post mitotic cnidocytes is sufficient for cnidocyte development, revealing that *NvLsd1* has a cell autonomous role and is not required for the initial specification of this cell type. This combined paradigm of an increase of Lsd1 levels as differentiation progresses and a requirement during late stages is also seen in certain cell types in vertebrates such as rod photoreceptors (32). The fact that the NvLsd1^K644A/A520E^ mutant is not capable of rescuing the cnidocyte phenotype suggests that catalytic function may be required for the role of NvLsd1 in cnidocytes. We cannot exclude, however, that this mutant form has also affects protein-protein interactions of NvLsd1 as has been shown for other LSD1 mutations (76).

The level of NvLsd1 is also higher in *NvElav1:*:mOrange^+^ and *NvFoxQ2d:*:mOrange^+^ neurons. We were, however, unable to pinpoint a morphological phenotype in these cell types upon mutation of *NvLsd1*. While this is in line with the small number of genes differentially expressed in mutants and upregulated in *NvElav1*::mOrange^+^ cells, we cannot rule out that morphological or functional alterations will become detectable with more sophisticated experimental tools.

In conclusion, we have shown that NvLsd1, a ubiquitously expressed chromatin modifier, is developmentally regulated in *Nematostella* and that it is required cell autonomously for the differentiation of cnidocytes. These observations allow us to speculate that developmental regulation of chromatin modifiers and their integration into cell differentiation programs arose early in animal evolution and thus constitute an ancient feature of metazoan development.

## Materials and Methods

### Animal care and maintenance

*Nematostella* were maintained at 18-19°C in 1/3 filtered sea water [*Nematostella* media (NM)], and spawned as described previously (77). Fertilized eggs were removed from their jelly packages by incubating in 3% cysteine in NM for 20 minutes followed by extensive washing in NM. Embryos were reared at 21°C and were fixed at 4 (cleavage stage), 8 (early blastula), 12 (mid blastula), 20 (early gastrula), 30 (late gastrula), 48 (early planula), 72 (mid planula), 96 (late planula), 120 (tentacle bud) hours post fertilization or at 13 days (primary polyp).

### Immunofluorescence (IF)

Animals older than 72 hours were anesthetized with MgCl2 and then killed quickly by adding a small volume (20-30 μl/ml) of 37% formaldehyde directly into the media. They were then fixed in ice cold 3.7% formaldehyde in PBTx [PBS(Phosphate Buffered Saline) + 0.2% Triton X-100] for 30-60 minutes (when staining for NvLsd1-GFP short fixations yield better staining) or for > 60 minutes or overnight (o/n) (for all other antibodies) at 4°C. Samples were washed >4 times in PBTx at RT, blocked in Block (3% BSA / 5% Goat serum in PBTx) for > 1 hour at RT and incubated in 1° antibody diluted in Block o/n at 4°C. Samples were then washed extensively in PBTx (> 5 washes for 2 hours or more) at RT, blocked for 1 hour at RT in Block and incubated o/n or over the weekend in 2° antibody diluted in Block at 4°C. If Phalloidin staining was performed, Alexa Fluor™ 488 or 633 Phalloidin (Thermo Fisher Scientific, A12379/ A22284) was added here at 1:50-1:100. Samples were then incubated in Hoechst 33342 (Thermo Fisher Scientific, 62249) at 1:100 in PBTx for 1 hour at RT followed by extensive washing in PBTx (> 5 washes for 2 hours or more). Animals were mounted in ProLong™ Gold Antifade Mountant with DAPI (Thermo Fisher Scientific, P36935) and imaged on a Leica SP5 confocal microscope. Antibodies and dilutions are listed in Table S1.

### DAPI staining for cnidocysts and counting

DAPI staining for cnidocysts was performed as previously published (60, 73) with slight modifications. Animals were processed as for IF with the addition of 10 mM EDTA to all solutions. Following the final PBTx wash, the samples were washed twice with MilliQ H_2_O and then incubated in 200 μg/ml DAPI in milliQ H_2_O o/n at RT. The samples were then washed once with MilliQ H_2_O, twice with PBTx with 10mM EDTA and mounted and imaged as for IF.

For counting of cnidocysts, late planula were imaged on a Leica SP5 confocal microscope and a maximum intensity projection of one half on the planula was generated using Fiji (78). The number of DAPI^+^ cnidocysts was then counted in a 100 μM square located in the middle of the planula. At least 10 planula were used per condition/ replicate. The experiment was repeated 3 times and the data shown is the results of one replicate which is representative of all. Statistical significance was assessed using a Student’s t-Test (2 tailed, equal variance).

### EdU labelling

Animals used in Edu labelling experiments were incubated in 10 mM EdU in NM for the desired time and then treated for IF as described. After the final set of PBTx washes EdU incorporation was visualized using the Click-iT™ EdU Imaging Kit with Alexa Fluor™ 488 or 647 (Thermo Fisher Scientific, C10337/C10337) following the manufacturer’s protocol. Samples were mounted and imaged as for IF.

### Flow cytometry and FACS

Flow cytometry and FACS sorting were performed as previously described (79). Briefly, animals were dissociated in 0.25% Trypsin (Gibco, 27250018) in Ca-Mg-free *Nematostella* media (CMFNM) (154 mM NaCl, 3.6 mM KCl, 2.4 mM Na_2_SO_4_, 0.7 mM NaHCO_3_) supplemented with 6.6 mM EDTA, pH 7.6-7.8. Cells were centrifuged at 800 g for 10 minutes, resuspended in ice cold 0.5% BSA in CMFNM (pH 7.6-7.8), filtered through a 40 μM filter and stained with Hoechst 33342 (Thermo Fisher Scientific, 62249) at a 60 μg/ml at RT for 30 minutes. Samples were then diluted 1:1 with ice cold 0.5% BSA/CMFNM and stained with 60 μl/ml 7-AAD (BD, 559925) for > 20 minutes on ice. Flow cytometry was performed on a BD Fortessa and sorting was carried out on a BD FACSAria II with a 100 μm nozzle. Data was analyzed using FlowJo v10.7.

### RNA extraction, library preparation and sequencing

For the RNAseq of *NvNcol3*::mOrange2^+^ cells, RNA extraction was performed as previously published (79). Cells were sorted directly into 0.5% BSA/ CMFNM at 4°C and centrifuged at 800 g for 10 minutes at 4°C. Most of the liquid was removed and 3 volumes of TRIzol LS reagent (Invitrogen, 10296028) was added. Samples were vortexed extensively and incubated at RT for 5 minutes before being flash frozen and stored at −80°C. The samples were processed using Direct-zol RNA MicroPrep columns (Zymo Research, R2060) including on column DNase digestion. cDNA was prepared from 400 pg of total RNA using the Smart-Seq 2 method with 16 pre-amplification PCR cycles (80). NGS libraries were prepared using the home-made tagmentation-based method (81). Briefly, 125 ng of cDNA was tagmented using home-made Tn5 loaded with annealed linker oligonucleotides for 3 minutes at 55°C. Reactions were inactivated by adding 1.25 ml of 0.2% SDS and incubation for 5 minutes at RT. Indexing and amplification was done using the KAPA HiFi HotStart PCR kit (Sigma-Aldrich, KK1508) with Index oligonucleotides (sequences were adapted from Illumina).

For RNAseq on late planula and primary polyp, 50 animals per sample were lysed in 500 μl TRIzol reagent (Invitrogen, 15596026) by vortexing extensively and incubated at RT for 5 minutes. 100 μl of chloroform was added and mixed vigorously and the aqueous component was isolated using MaXtract High Density tubes (Qiagen, 129046) using the manufacturers protocol. One volume of 100% Ethanol was added to the aqueous phase. This was then processed using an RNeasy Micro Kit (Qiagen, 74004) using the manufacturers protocol including on column DNase digestion using the RNase-Free DNase Set (Qiagen, 79254). Libraries were prepared using the NEBNext Ultra II Directional RNA Library Prep Kit for Illumina (E7760L), with following changes: 25ng RNA input, 1/100 adaptor dilution, 14 PCR cycles. Libraries were sequenced using a 75bp single end sequencing on a NextSeq500 machine (Illumina). RNA quality was assessed using an RNA 6000 Pico Kit (Aligent, 5067-1513) and concentration determined using the Qubit™ RNA HS assay kit (Invitrogen, Q32852).

### sgRNA synthesis

For the generation of the *NvLsd1* mutant allele the sgRNA was produced using a template generated by primer annealing. A PCR was set up containing 5 μl of each primer (100 mM), 2 μl dNTPs (10 mM each), 2 μl Q5 polymerase (NEB, M0491), 10 μl Q5 reaction buffer and 31 μl H2O with the following protocol: 98°C, 90 seconds; 55°C, 30 seconds; 72°C, 60 seconds. This was purified using a PCR clean up kit (Promega, A9281). The sgRNAs were synthesized using the MEGAscript™ T7 Transcription Kit (Invitrogen, AMB13345) including the DNase treatment.

For the generation of *NvLsd^GFP^* line, sgRNAs were produced using the EnGen^®^ sgRNA Synthesis Kit (NEB, E3322S). Primers are given in Table S2.

In both cases, sgRNAs were precipitated by adding 1:1 LiCl (7.5 M) (Invitrogen AM9480) and incubating at −20°C for 30 minutes followed by centrifugation at full speed at 4°C for 15 minutes and extensive EtOH washes. The concentration was calculated using a nanodrop.

### CRISPR-Cas9 injections and genotyping

*NvLsd1* mutants were produced similarly to previously published (82, 83). Eggs were injected with a mix containing sgRNA (130 ng/μl), Cas9 (PNA Bio, CP01) (500 ng/μl) and 1:4 Dextran, Alexa Fluor™ 568 (Invitrogen, D22912) (200 ng/μl in 1.1 M KCl) that was incubated at 37°C for 5-10 minutes prior to injection. Injected animals were raised to sexual maturity and crossed to wildtypes. F1 offspring were analyzed by sequencing in order to identify an F0 carrying the desired mutation. Individual F1s were placed in tubes, the NM removed and 100% EtOH added. After 5 minutes this was removed and the tubes were placed at 50°C for 45 minutes to allow the remaining EtOH to evaporate. 50 μl genomic extraction buffer (10 mM Tris pH8, 1 mM EDTA, 25 mM NaCl, 200 μg/μl ProteinaseK) was added to each and incubated at 50°C for 2 hours and 98°C for 15 minutes. 2 μl of this was used for PCR and sequencing. Once an F0 carrier was identfied, the remaining F1 offspring from that carrier were genotyped using a cut piece of tissue in order to generate a pool of F1 heterozygous animals.

For generating the *NvLsd1^GFP^* line, the repair template was produced by PCR using primers given in Table S2 and an in-house eGFP plasmid as template. The PCR product was gel extracted (Promega, A9281). Wildtype embryos were injected with a mix containing two sgRNAs (56.25 ng/μl each), repair template (25 ng/μl), Cas9 (PNA Bio, CP01) (500 ng/μl) and 1:4 Dextran, Alexa Fluor™ 568 (Invitrogen, D22912) (200 ng/μl in 1.1 M KCl) that was incubated at 37°C for 5-10 minutes prior to injection. Animals were screened for GFP fluorescence in the days following injection and GFP^+^ animals were grown to maturity and crossed to wildtypes in order to identify a carrier. A single F1 male was used to generate the line and all animals used in further analysis are offspring of this animal. The insertion was validated using PCR and western blotting (See Figure S2).

### PCR, cloning and sequencing

For generating cDNA for PCR, RNA was extracted as for the RNAseq of mutants. The SuperScript™ III first-strand synthesis system (Invitrogen, 18080051) was used to generate cDNA. The gDNA for PCR was extracted as for genotyping except from several pooled animals.

All PCRs were performed with Q5 polymerase and primers are listed in Table S2.

For cloning of the *NvLsd1-GFP* cDNA and genomic DNA fragments (Fig. S2) the fragments were cloned using the CloneJET PCR Cloning Kit (Thermo Fisher Scientific, K1231). For the cloning of the *NvLsd1* cDNA primers were designed using the gene model (Nve23413, JGI: v1g105193) and the amplified fragment was cloned into a pCR4 backbone using the NEBuilder^®^ HiFi DNA Assembly master mix (NEB, E2621). For sub-cloning into the *NvPOU4*::mCherry plasmid (52) both the backbone and the insert were amplified by PCR and assembled using the NEBuilder^®^ HiFi master mix. The *NvH2B* was amplified from primers designed against gene model Nve18479 (Genebank: XM_001635370.2). To generate the NvLsd1^K644A/A520E^ mutations we used synthesized fragments of *NvLsd1* continuing the mutations. We then amplified the *NvPOU4*::NvLsd1-mCherry plasmid to remove the corresponding part of *NvLsd1* and inserted the fragment using the NEBuilder^®^ HiFi master mix.

Plasmids were sequenced using either BigDye™ Terminator v3.1 sequencing kit (Applied Biosystems, 4337458) and analyzed in house or samples were sequenced commercially using the GENEWIZ sanger sequencing service.

### Bioinformatic analysis

The quality of raw RNA-seq reads was initially assessed with FastQC software v.0.11.8 (84) and filtering was performed with fastp v.0.20.0 (85) in default settings. Reads which passed quality control were mapped with STAR aligner v.2.7.3a (86) in default settings to *N.vectensis* genome (https://mycocosm.jgi.doe.gov/Nemve1/Nemve1.home.html) (87) using NVE gene models (https://figshare.com/articles/Nematostella_vectensis_transcriptome_and_gene_models_v2_0/807696). Downstream analysis was performed using R v.4.0.2 (88) and Bioconductor packages (https://www.bioconductor.org). Briefly, aligned reads were counted with ‘summarizeOverlaps’ function from the package ‘Genomic Alignments’ v.1.24.0 (89) and genes with less than 10 counts in at least 4 biological replicates for one condition were filtered out. Differential expression analysis was performed using ‘DESeq2’ package v.1.28.1(90). Overlap between up- and/or down-regulated genes across different conditions was assessed with ‘GeneOverlaps’ package v.1.23.0 (91).

### Analysis of Lsd1 mutant survival, metamorphosis and growth

In order to assess the survival of *NvLsd1* mutant embryos Lsd1^GFP/-^ animals were in-crossed. At 48 hours, ~100 embryos were sorted as having “normal” morphology independent of whether they were GFP^+^. At 96 hours these animals were genotyped based on whether they were GFP^+^ (control) or GFP^−^ (mutant). These same animals were separated into 2 dishes based on their genotype and grown to primary polyp. The number of these animals to have properly developed was then counted at this stage and is shown as a percentage of the number of animals present at 4 days. These were then imaged on a Nikon SMZ18 with a Nikon DS-Qi2 camera, using the same magnification and settings and the length of each animal was determined using the measure tool in Fiji (78). Length was normalized by dividing by the mean length of the control animals. Statistical significance was assessed using the Student’s t-Test (2 tailed, equal variance). Levene’s Test was used to determine equal variance and normalily was tested using the Kolmogorov-Smirnov Test.

### Transgenesis

In order to generate F0 mosaic transgenics we used I-Sce1 mediated transgenesis as previously described (92) with minor modifications. Eggs were injected with a mix containing: plasmid DNA (10ng/ul), ISce1 (1U/ul) (NEB, R0694), Dextran Alexa Fluor™ 568 (100ng/ul), CutSmart buffer (1x). The mix was incubated for 30 minutes at 37°C before injection.

### Data availability

All sequencing data will be made freely available upon publication and is available on request.

## Acknowledgements

We thank the members of the Rentzsch lab for support and discussions throughout the project, Eilen Myrvold and Lavina Jubek for excellent care of animals in the Sars Centre *Nematostella* facility, Yehu Moran for providing both the *NvNcol3*:mOrange2 and *NvPTx1*:mOrange2 transgenic lines, Suat Özbek for the NvNcol3 antibody, Brith Bergum for performing cell sorting and for training and support for the flow cytometry experiments, Pawel Burkhardt and Tarja Hoffmeyer for sharing reagents and providing advice on western blotting, Marios Chatzigeorgiou for helpful discussions and advice and Gemma Richards for initial in situ hybridizations. We are grateful to Eric Röttinger and Jake Warner for help and advice in generating the NvLsd1^GFP^ knock-in line. Cell sorting and flow cytometry was performed at the Flow Cytometry Core Facility, Department of Clinical Science, University of Bergen. Sequencing was performed at EMBL Genecore. Research in FRs lab was funded by a grant from the University of Bergen and the Research Council of Norway (251185/F20) and by the Sars Centre core budget.

## Author Contributions

J.M.G designed and performed the experimental work, analyzed the data, conceptualized the study, generated the figures, and wrote the manuscript. I.U.K. analyzed and visualized the transcriptome data. F.R. conceptualized and supervised the study and wrote the manuscript. All authors edited the manuscript.

