## Supplementary material for "Histone demethylase Lsd1 is required for the differentiation of neural cells in the cnidarian *Nematostella vectensis*": Supplemetary material

This PDF includes:

Supplementary Material and Methods
Tables S1 and S2
Figures S1- S8
Supplementary references

### Supplementary Materials and Methods

**RNA *In-situ* hybridization.** Animals were fixed in ice cold 0.2% glutaraldehyde/3.7% formaldehyde in NM for 1.5 minutes followed by 1 hour at 4°C in 3.7% formaldehyde in PBT (PBS + 0.1% Tween 20). Animals were washed several times in PBT at room temperature (RT) and dehydrated through a series of methanol washes and stored in 100% methanol at -20°C. Probes were generated with T3 RNA polymerase (Roche, 11031163001) and DIG RNA labelling mix (Roche 11277073910). *In situ* hybridization was performed as previously described (1, 2). Samples were imaged on a Nikon Eclipse E800 compound microscope with a Nikon Digital Sight DSU3 camera.

**Western blotting.** Protein extraction was performed on $Lsd^{GFP}$ or wild-type late planula. Animals were placed in RIPA buffer (150mM NaCl, 50mM Tris pH8, 1% NP40, 0.5% DOC, 0.1% SDS) supplemented with cOmplete EDTA-free Protease Inhibitor Cocktail (Roche, 4693159001) and homogenized by passing through a 27G needle. Samples were incubated on ice for 30 minutes and mixed by passing through the needle every 5 minutes and centrifuged at full speed for 15 minutes @ 4°C. The supernatant was kept and the protein concentration quantified using the Qubit™ Protein Assay (Invitrogen, Q33212). 30 µg of protein was used per lane, mixed 1:1 with 2X Laemmli sample buffer (0.1M TrisHCl pH 6.8, 2% SDS, 20% Glycerol, 4% β-mercaptoethanol 0.02% Bromophenol blue) and boiled for 5 minutes before loading. PageRuler™ Plus prestained protein ladder, 10 to 250 kDa (Thermo Scientific, 26619) was used. SDS PAGE was performed using 7.5% Mini-PROTEAN® TGX™ precast protein gels (BIO-RAD, 4561023) run in running buffer (25mM Tris, 192mM Glycine, 0.1% SDS) at 100 V for ~120 minutes. Transfer was performed using Trans-Blot Turbo Mini 0.2 µm PVDF Transfer Pack (BIO-RAD, 1704156) on a Trans-Blot Turbo transfer system (BIO-RAD) using the high molecular weight program. After transfer the membrane was washed in PBT (PBS + 0.1% Tween) several times and blocked with 5% milk powder in PBT (MPBT) at RT for 1 hour. The blots were incubated o/n at 4°C in 1º antibody (Ab290) in MPBT. The membranes were then washed several times in PBT and incubate in 2º antibody in MPBT at RT for 1 hour. Membranes were then washed several times in TBT and the signal was revealed using Clarify ECL substrate (BIO-RAD, 1705060) and imaged on a ChemiDoc XRS+ (BIO-RAD). The blots were then washed in PBT and blocked again in 5% MPBT for 1 hour at RT. They were then incubated o/n at 4°C with anti-actin antibody and processed as for the first antibody. Antibodies are in Table S2.<u>Table S1. List of antibodies used in this study</u>

#### Supplementary Figures and Tables

Table S1. List of antibodies used in this study

| Name | Company | Catalogue number | Concentration (IF) | Concentration (Western) |
|---|---|---|---|---|
| Rabbit anti-DsRed | Clontech | 632496 | 1:100 | |
| Mouse anti-mCherry | Clontech | 632543 | 1:100 | |
| Rabbit anti-GFP | Abcam | Ab290 | 1:200 | 1:20,000 |
| Mouse anti-GFP | Abcam | Ab1218 | 1:200 | |
| Goat anti-rabbit Alexa 488 | Life Technologies | A11008 | 1:250 | |
| Goat anti-rabbit Alexa 568 | Life Technologies | A11011 | 1:250 | |
| Goat anti-mouse Alexa 488 | Life Technologies | A11001 | 1:250 | |
| Goat anti-mouse Alexa 568 | Life Technologies | A11004 | 1:250 | |
| Rabbit anti-Actin | Abcam | A5060 | | 1:1000 |
| Goat Anti-Rabbit (HRP) | Abcam | Ab97051 | | 1:10,000 |

#### <u>Table S2. List of primers used in this study</u>

| Name | Use | Sequence |
|---|---|---|
| *NvLsd1*_sgRNA1_Fw | Generating the template for the sgRNA used to generate the *NvLsd1* mutant | 5’ TAATACGACTCACTATAGGGTTTTGGCAACCTTAATAGTTT TAGAGCTAGAA 3’ |
| *sgRNA*_Rv | | 5’AAAAGCACCGACTCGGTGCCACTTTTTCAAGTTGATAACGG ACTAGCCTTATTTTAACTTGCTATTTCTAGCTCTAAAAC 3’ |
| *NvLsd1*_K/in_sgRNA1 | Generating the sgRNAs used to generate the NvLsd1$^{GFP}$ animals | 5’ TTCTAATACGACTCACTATAGGCATATATCTAGCGAGGTGTT TTAGAGCTAGA 3’ |
| *NvLsd1*_K/in_sgRNA2 | | 5’ TTCTAATACGACTCACTATAGGATATATCTAGCGAGGTTGGTT TTAGAGCTAGA 3’ |
| *NvLsd1*_Mut_Seq_Fw | PCR for sequencing the *NvLsd1* mutants | 5’ GCCGACGCCGTACTC 3’ |
| *NvLsd1*_Mut_Seq_Rv | | 5’ GCCTATAATGTACGAGCTGGTTTTGGC 3’ |
| Repair_Template_Fw | Generating the repair template for the Knock-in | 5’ CGAAGTCGGGCGAGTGAGCCCACCCATGCCCCAACCTCGC GGCGGAGGAGGCTCAGG 3’ |
| Repair_Template_Rv | | 5’ GTTCGTCCGTGATACGCAGGGTCGGTGCAGCATATATCTAC TTGTCGTCATCGTCTTTGTAGTCCTTGTACAGCTCGTCCATGC3’ |
| NvLsd1_F1 | PCR to confirm the GFP insertion in NvLsd1$^{GFP}$ animals (See Fig. S3) | 5’ CGGCTAACGTGGCACCAG 3’ |
| NvLsd1_R1 | | 5’GACAACTATTTCGTAGGCTCAAATTTCGCC 3’ |
| NvLsd1_F2 | To amplify the Lsd1$^{GFP}$ cDNA( See Fig. S3) | 5’ ATGTCGATTCCACCATATCAAACTC 3’ |
| NvLsd1_R2 | | 5’ CTAGCGAGGTTGGGGC 3’ |
| NvLsd1_cDNA_Fw | To amplify *NvLsd1* coding sequence | 5’ ATGTCGATTCCACCATATCAAACTC 3’ |
| NvLsd1_cDNA_Rv | | 5’ CAGTTTCAGAGATTACAAGTTGAAAGGTTTTC 3’ |
| NvLsd1_ATG_Fw_1HA | Amplify *NvLsd1* and add the 2HA tags | 5’ GACTACGCAGGTTACCCTTACGATGTTCCCGACTACGCAGG TTCCTCGATTCCACCATATCAAACTCCAATTCTACC 3’ |
| NvLsd1_ATG_Fw_2HA | | 5’TAGATATCAGTTGGTGCTGAGTGCTCCACGATGTATCCCTAT GATGTTCCAGACTACGCAGGTTACCCTTACGATG 3’ |
| NvLsd1_noSTOP_Rv | | 5’ GCGAGGTTGGGGCATGG 3’ |
| *NvPOU4*:mCherry_Fw1 | Amplify the *NvPOU4*:mCherry backbone with overhangs for *NvLsd1* | 5’ GCCCACCCATGCCCCAACCTCGCGCGGGCGGAGGTGGCA GCGTGAGCAAGGGCGAGGAAG 3’ |
| *NvPOU4*:mCherry_Rv1 | | 5’ ATCGTAAGGGTAACCTGCGTAGTCTGGAACATCATAGGGAT ACATCGTGGAGCACTCAGCACCAAC 3’ |
| NvH2B_Fwd | Amplify *NvH2B* | 5’ ATGCCTGCCAAGAAAAGAGCTCC3’ |
| NvH2B_NoSTOP_Rv | | 5’ TCCAGTGCTCGCTGAGTATTTGG 3’ |
| *NvPOU4*:mCherry_Fw2 | Amplify the *NvPOU4*:mCherry backbone with overhangs for *NvH2B* | 5’GCCGTCGCCAAATACTCAGCGAGCACTGGAGCGGGCGGCG GCGGCAGC3’ |
| *NvPOU4*:mCherry_Rv2 | | 5’CCCGGCAGGAGCTCTTTTCTTGGCAGGCATCGTGGAGCACT CAGCACCAAC 3’ |
| *NvLsd1*_Internal_Fw1 | Amplify the *NvPOU4*:NvLsd1-mCherry plasmid to insert the K644A mutation | 5’ GCACACACGTGGGGAGTTGTTTC 3’ |
| *NvLsd1*_Internal_Rv1 | | 5’ GGAACTGCACAGCGGCAG 3’ |
| *NvLsd1*_Internal_Fw2 | Amplify the *NvPOU4*:NvLsd1-mCherry plasmid to insert the A520E mutation | 5’ CAGTGAGACATGTGCGCTACAGTAG 3’ |
| *NvLsd1*_Internal_Rv2 | | 5’ CCAGCTCTTCTAATCTCTCGTCCAGC 3’ |

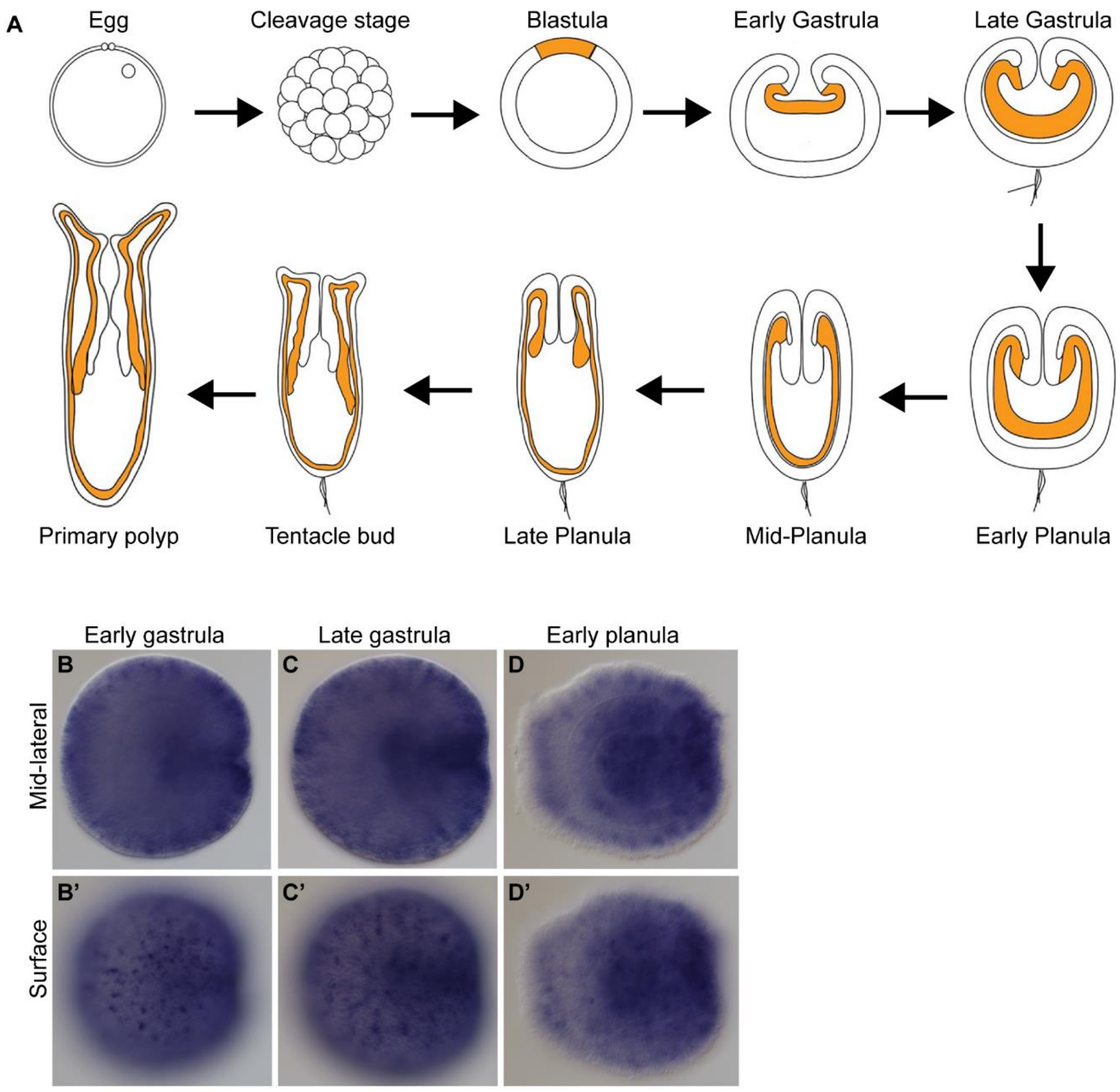


**Figure S1** ***Nematostella*** **life cycle and** ***NvLsd1*** **expression**. (**A**) Schematic representation of *Nematostella* developmental stages shown throughout the paper. The endomesoderm is shown in orange. Adapted from (3). (**B**) RNA in-situ hybridization for *NvLsd1* during embryogenesis. Stage is shown on top. (**B-D**) show mid-lateral views and (B'-D') show surface views.

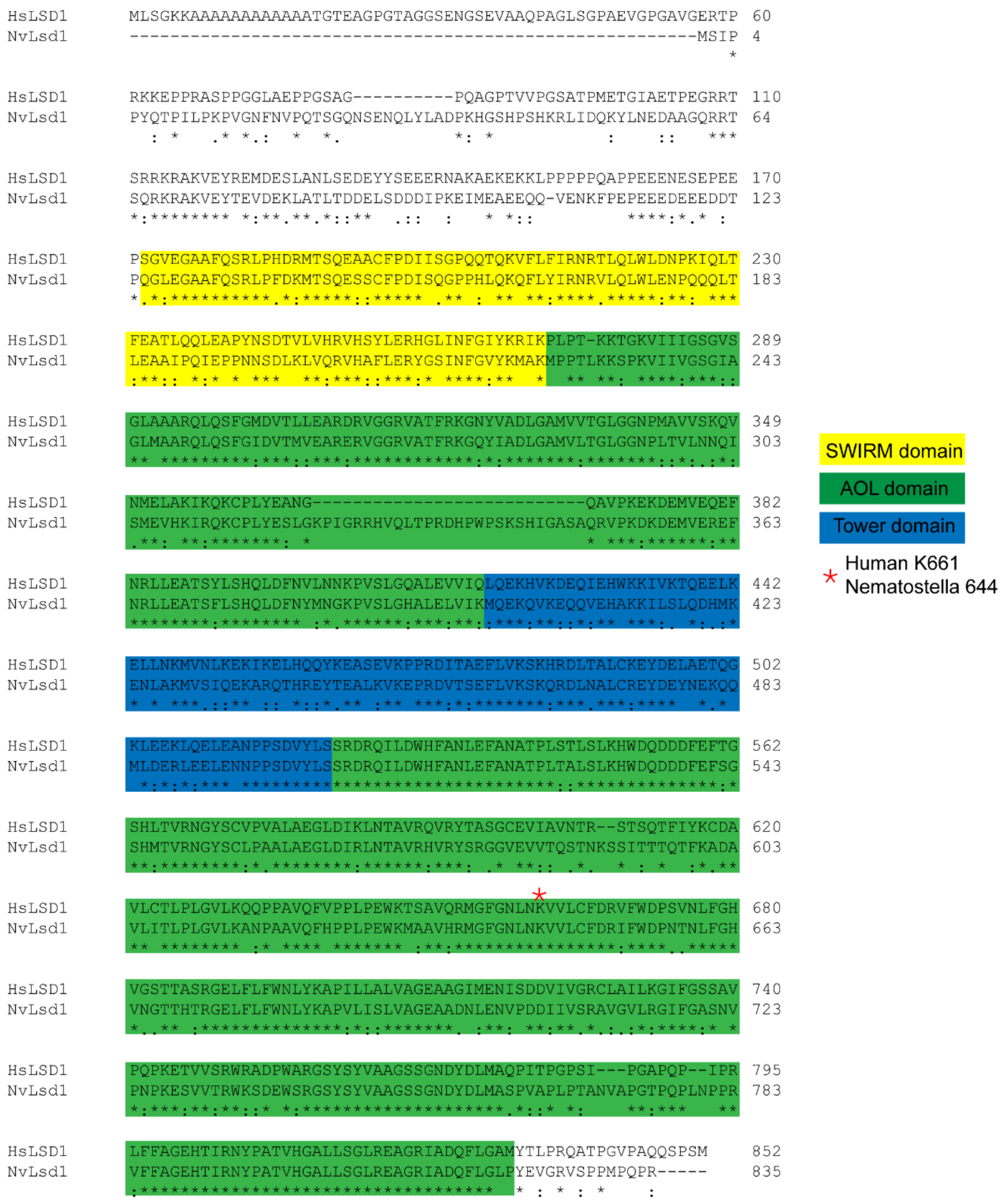


**Figure S2: Alignment of human and *Nematostella* Lsd1 sequences**
Alignment of the full-length NvLsd1 sequence with the human LSD1. The SWIRM domain is highlighted in yellow, the Amine Oxidase-like (AOL) domain in green and the Tower domain in blue. The position of human lysine 661/ *Nematostella* lysine 644 is marked with a red asterisk. The alignment was performed using Clustal Omega (4).

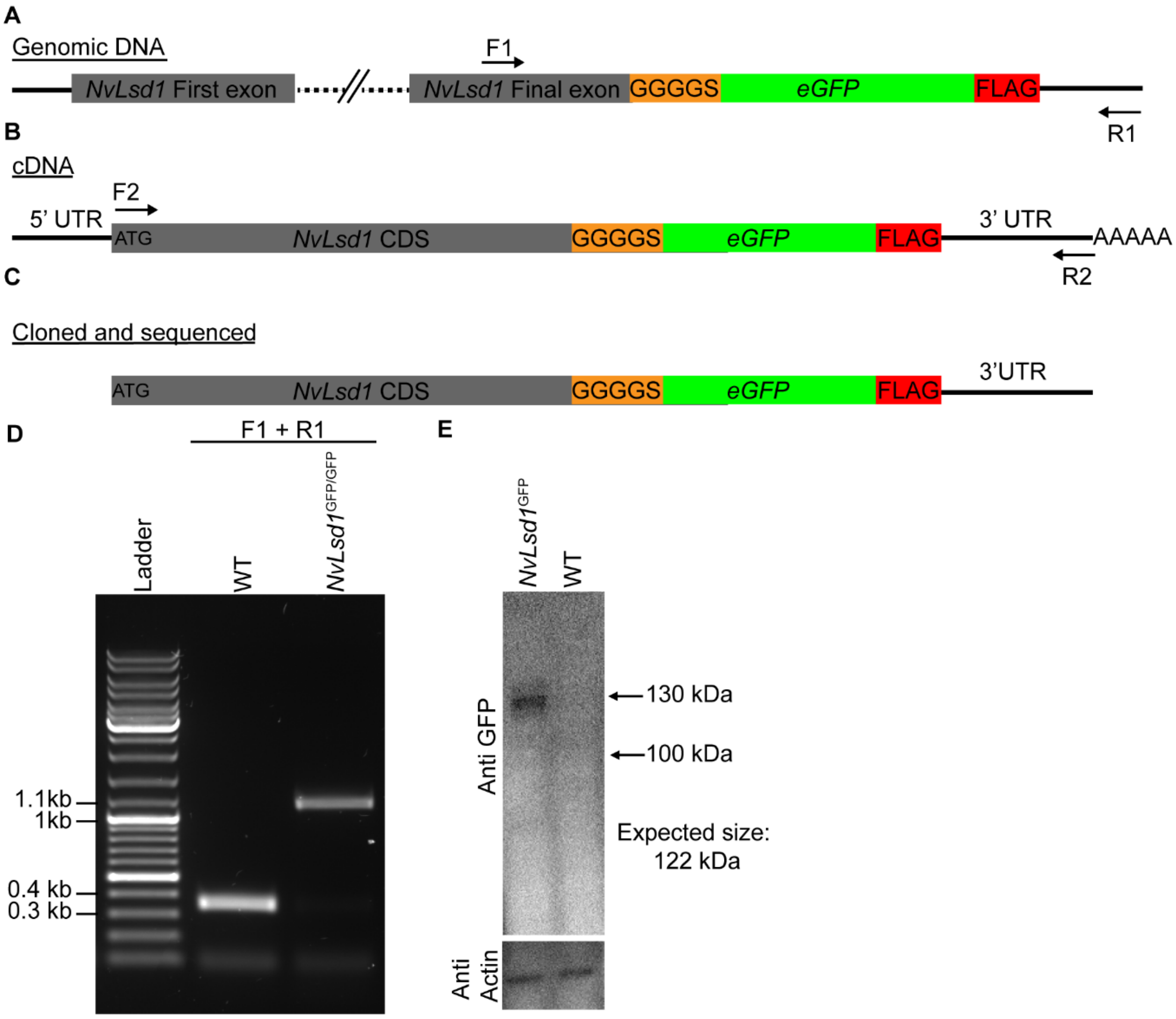


**Figure S3: Confirmation of *GFP* knock-in into the *NvLsd1* locus.**
(**A**) Schematic showing the insertion of GFP into the genome in frame with *NvLsd1* and indicating the position for the primers used to verify the insertion. (**B**) Schematic of the predicted full length *NvLsd1$^{GFP}$* cDNA with the inserted sequences. F3 and R3 indicate the primers used to amplify and clone the full length coding sequence and 5'UTR from *NvLsd1$^{GFP}$* animals. (**C**) Schematic of the portion of *NvLsd1$^{GFP}$* cDNA which was cloned and sequenced. (**D**) Agarose gel of a PCR on genomic DNA from WT and homozygous *NvLsd1$^{GFP}$* animals showing the insertion of a single copy of GFP into the *NvLsd1* locus. The primers used are indicated on top. The expected size of the band from wildtype (WT) is 336 bps and from *NvLsd1$^{GFP}$* animals is 1089 bps. The band in the *NvLsd1$^{GFP}$* lane was excised and sequenced to confirm it is the correct sequence. (**E**) Western blot on protein from *NvLsd1$^{GFP}$* and wild-type (WT) animals. The blot was probed for GFP (shown on top) and Actin (shown on the bottom). Sizes on the right indicate the positions of the ladder. The expected size of the NvLsd1-GFP fusion protein is 122 kDa,

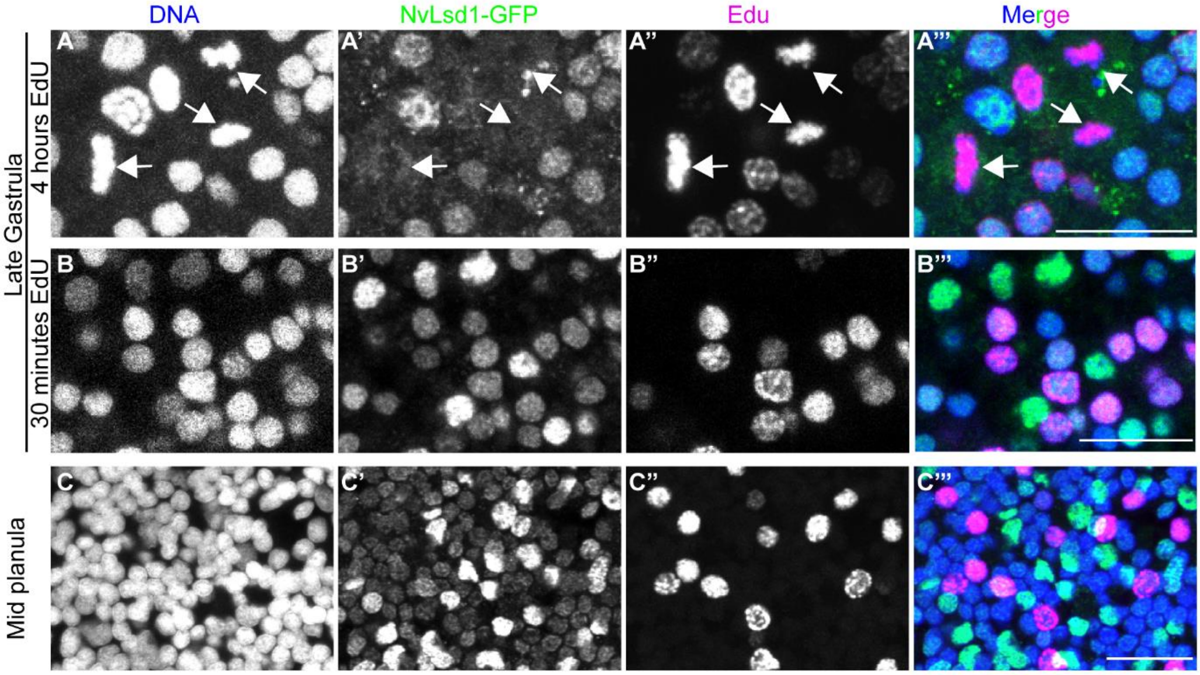


**Figure S4: Edu labelling in *NvLsd1*$^{GFP}$ animals.**

(**A-C**) Confocal images showing close up of the ectoderm of a late gastrula incubated with Edu for 4 hours (**A**), late gastrula inbuated with Edu for 30 minutes (**B**) and mid-planula incubated with EdU for 30 minutes (**C**). Stage is shown on the left. Arrows in (**A**) point to mitotic nuclei. ClickIT-Edu is shown in magenta, Lsd1-GFP in green and DNA in blue. Scale bars: 10 µm

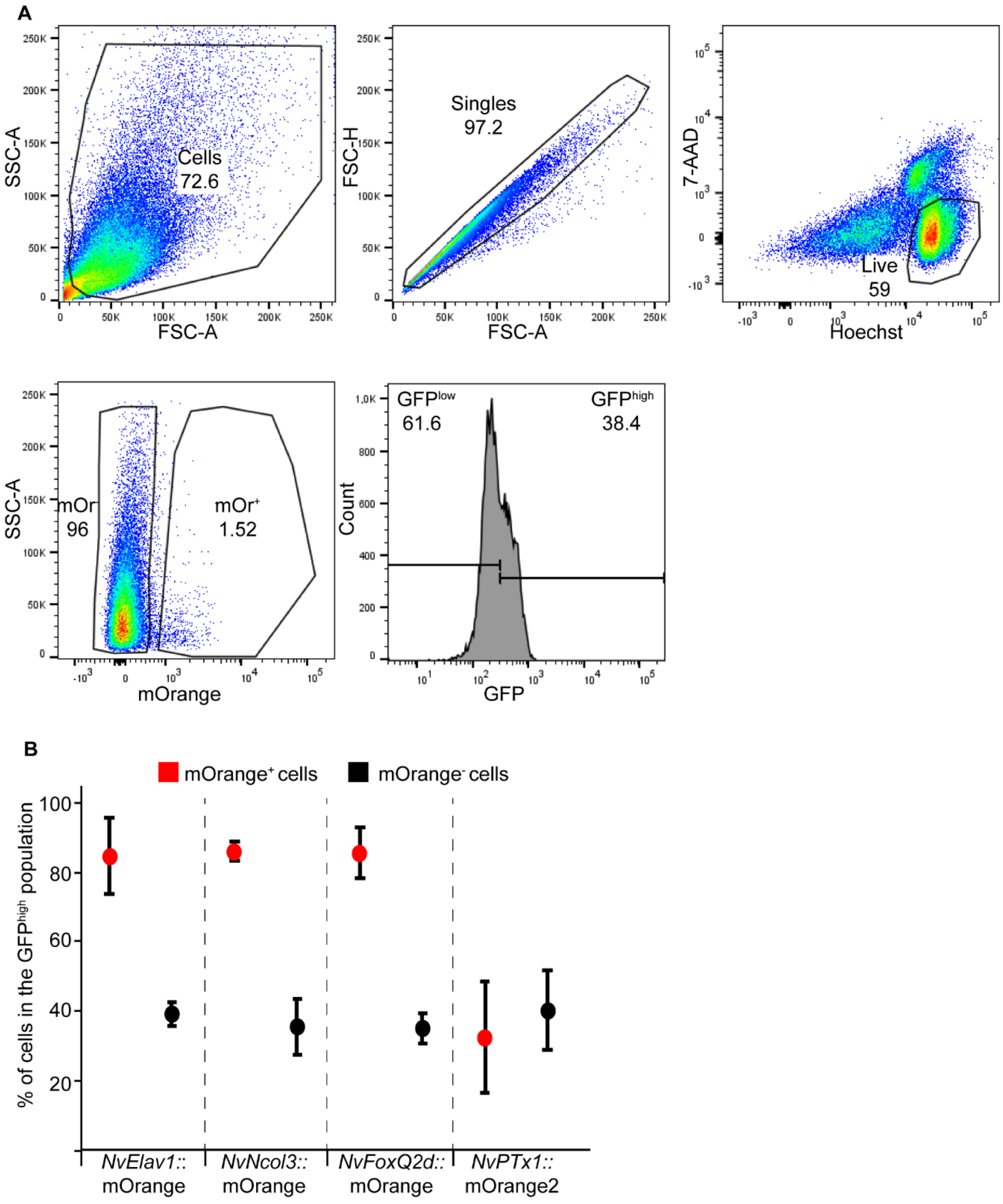


**Figure S5: Flow cytometry analysis of *NvLsd*$^{GFP}$ animals**

(**A**) FlowJo plots showing an example of the gating strategy used to analyze the flow cytometry data. (**B**) Plots showing the percentage of mOrange$^{+}$ (red) and mOrange$^{-}$ (black) cells that fall within the GFP$^{high}$ gate. The data shows the average of 3 independent replicated and the error bars show the standard deviation.

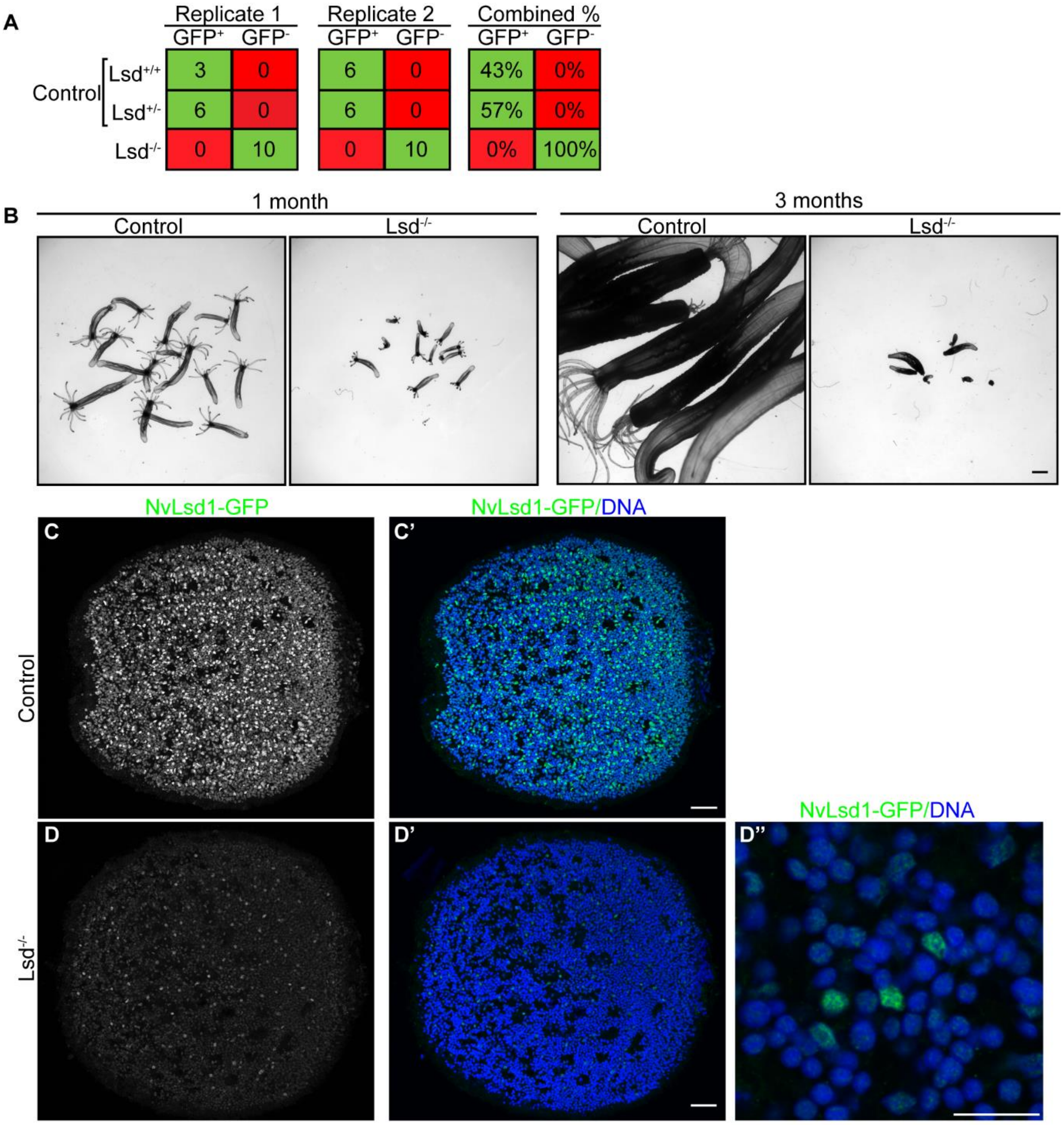


**Figure S6: Further characterization of *NvLsd1* mutants**

(**A**) Genotyping of GFP$^+$ and GFP$^-$ animals from the crosses shown in Figure 3A. (**B**) Images of animals shown in Figure 3F-I after 1 month and 3 months. (**C, D**) Anti-GFP immunostaining in Control or *NvLsd1*$^{-/-}$ mid-planula. D'' shows a close up taken at a higher detector threshold. Scale bars: 1 mm (B), 25 µm (C–D') and 10 µm (D'').

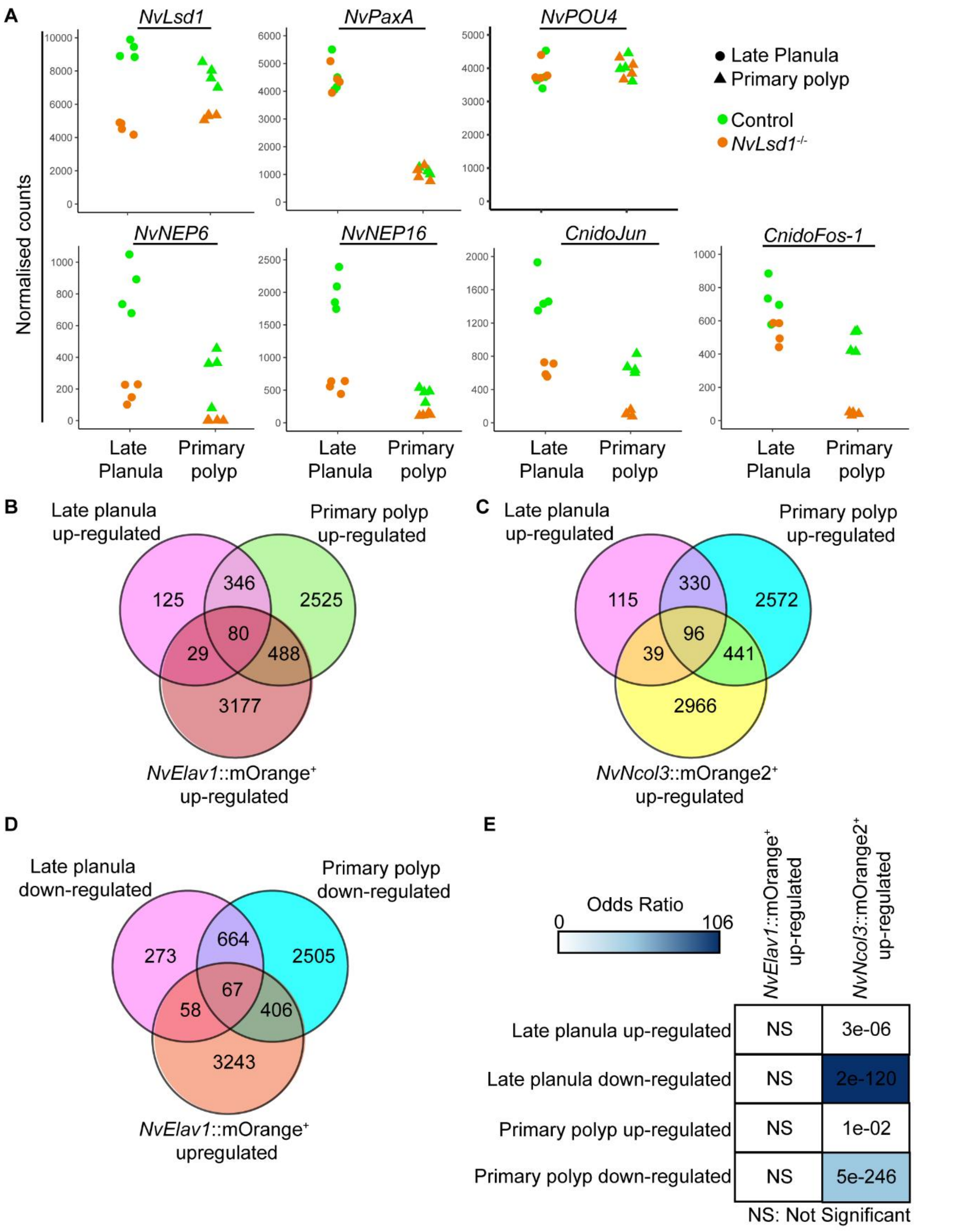


**Figure S7. Comparison of differentially expressed genes in *NvLsd1* mutant with neural transcriptomes.**

(**A**) Plots showing normalized count data for selected genes in *NvLsd1* mutants. Control samples are shown in green and mutants in orange. Late planula samples are represented by circles and primary polyp samples are shown as triangles. (**B**, **C**) Venn diagram comparing genes up-regulated in *NvLsd1*$^{-/-}$ late planula and primary polyp with genes up-regulated in *NvElav1*::mOrange$^{+}$ (B) and *NvNcol3*::mOrange2$^{+}$ (C) cells. (**D**) Venn diagram comparing genes down-regulated in *NvLsd1*$^{-/-}$ late planula and primary polyp with genes upregulated in *NvElav1*::mOrange$^{+}$ cells. (**E**) Comparison of the overlap between up- and down-regulated genes in *NvLsd1*$^{-/-}$ late planula and primary polyps with those up-regulated in *NvNcol3*::mOrange2$^{+}$ and *NvElav1*::mOrange$^{+}$ cells using the GeneOverlap R package using genes with a log2 fold change threshold of 1. The strength of the blue color indicates the odds ratio and numbers indicated the p-value, calculated using Fisher's exact test.

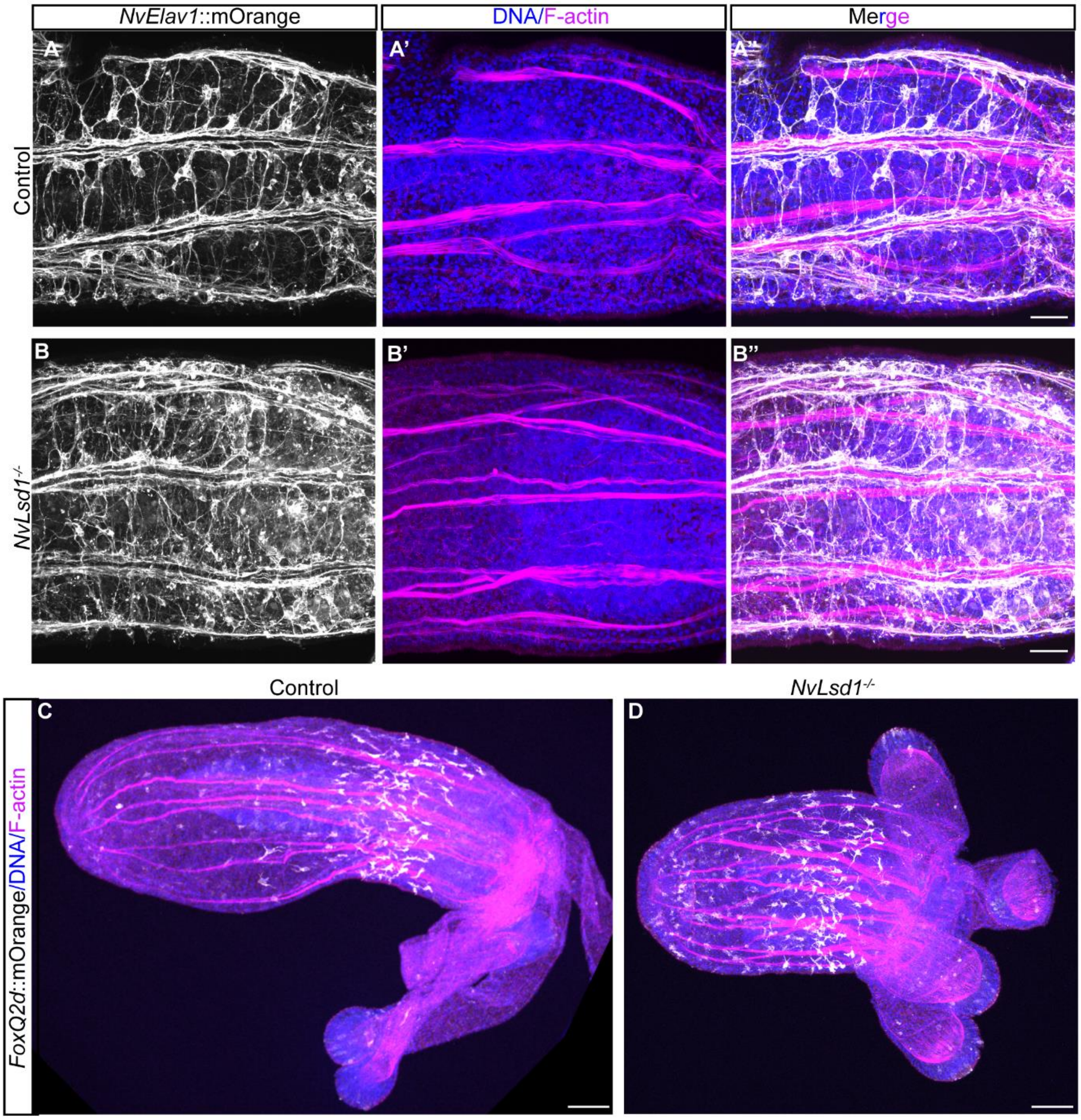


**Figure S8: Analysis of *NvElav1*::mOrange and *NvFoxQ2d*::mOrange transgenes in *NvLsd1* mutants.**

(**A-D**) Confocal images of immunofluorescence staining on control and *NvLsd1*$^{-/-}$ primary polyps in the background of the *NvElav1*::mOrange (**A, B**) *NvFoxQ2d*::mOrange (**C**, **D**) transgenes. (**A, B**) show close ups of the endoderm and (**C, D**) show full primary polyps. mOrange in shown in gray, DNA in blue and F-actin in magenta. Scale bars: 20 µm (A-B) and 50 µm (C, D).
